# Spatial evidence of cryptic methane cycling and methylotrophic metabolisms along a land-ocean transect in a southern California salt marsh

**DOI:** 10.1101/2024.07.16.603764

**Authors:** Sebastian J.E. Krause, Rebecca Wipfler, Jiarui Liu, David J. Yousavich, DeMarcus Robinson, David W. Hoyt, Victoria J. Orphan, Tina Treude

**Affiliations:** Department of Earth Planetary and Space Sciences, University of California, Los Angeles, CA 90095, USA; Department of Atmospheric and Ocean Sciences, University of California, Los Angeles, CA 90095, USA; Division of Geological and Planetary Sciences, California Institute of Technology, Pasadena, CA 91125, USA; Pacific Northwest National Laboratory Environmental and Molecular Sciences Division, Richland, WA 99352, USA

**Author notes:** Earth Research Institute, 6832 Ellison Hall, University of California Santa Barbara, Santa Barbara, CA 93106-3060, USA.

**Keywords:** Sulfate reduction, Iron reduction, Anaerobic oxidation of methane, Methanogenesis, Mono-methylamine, Sulfate-reducing bacteria, Methanogenic archaea, Methanotrophic archaea

## Abstract

Methylotrophic methanogenesis in the sulfate reduction zone of coastal and marine sediments couples with anaerobic methane oxidation (AOM), forming the cryptic methane cycle. This study provides evidence of cryptic methane cycling in the sulfate-reducing zone across a land-ocean transect of four stations—two brackish, one marine, and one hypersaline—within the Carpinteria Salt Marsh Reserve (CSMR), Southern California, USA. The top 20 cm of sediment from the transect underwent geochemical and molecular (16S rRNA) analyses, in-vitro methanogenesis incubations, and radiotracer incubations using ^35^S-SO_4_, ^14^C-mono-methylamine, and ^14^C-CH^4^. Sediment methane concentrations were consistently low (3 to 28 µM) except at the marine station, where they increased with depth (max 665 µM). Methanogenesis from mono-methylamine was detected throughout the sediment at all stations with estimated rates ranging between 0.14 and 3.8 nmol cm^−3^ d^−1^. 16S rRNA analysis identified methanogenic archaea capable of producing methane from methylamines in sediment where methanogenesis was found to be active. Metabolomic analysis of porewater showed mono-methylamine was mostly undetectable (<3 µM) or present in trace amounts (<10 µM) suggesting rapid metabolic turnover. In-vitro methanogenesis incubations showed no linear methane buildup, suggesting a process limiting methane emissions. AOM activity, measured with ^14^C-CH_4_, overlapped with methanogenesis from mono-methylamine activity at all stations, with rates ranging from 0.03 to 19.4 nmol cm^−3^ d^−1^. Porewater geochemical analysis showed the CSMR sediments are rich in sulfate and iron. Porewater sulfate concentrations (9–91 mM) were non-limiting across the transect, which support members of sulfate-reducing bacteria and likely responsible for sulfate reduction activity (1.5–2,506 nmol cm^−3^ d^−1^) at all stations in the CSMR. Porewater sulfide and iron (II) profiles indicated that the sediment transitioned from a predominantly iron-reducing environment at the two brackish stations to a predominantly sulfate-reducing environment at the marine and hypersaline station. AOM activity was likely coupled to sulfate and possibly iron reduction, coinciding with the presence of anaerobic methanotrophs and bacteria involved in these reductions. 16S rRNA analysis identified anaerobic methanotrophs at the marine and hypersaline stations, where they coexisted with putative methanogens, suggesting both groups, or methanogens alone, may be involved in cryptic methane cycling, preventing significant methane buildup in the sulfate-reducing zone. Differences in rate constants from ^14^C radiotracer incubations suggest a non-methanogenic process oxidizing mono-methylamine to inorganic carbon, likely mediated by sulfate-reducing bacteria. Understanding the potential competition of sulfate reducers with methanogens for mono-methylamine needs further investigation as it might be another important process responsible for low methane emissions in salt marshes.

## Introduction

Methane is the simplest and most abundant organic molecule in the atmosphere and is about 25-30 times more potent than carbon dioxide as a greenhouse gas (IPCC, 2014). Since pre-industrial technological advancement, methane concentrations in the atmosphere have nearly tripled from 722 ppb to approx. 1900 ppb in 2022 (Wang et al., 2024). Saunois et al. (2020) reported that natural wetlands between the years 2008-2017 contributed on average 142 Tg CH_4_ yr^−1^ (range between 102-182 Tg CH_4_ yr^−1^) which makes natural wetlands the largest contributor of methane into the atmosphere. Natural wetlands are broadly characterized into freshwater and coastal wetlands. Both ecosystems contain organic-rich sediment that could sustain microbial methanogenesis. However, coastal wetlands, such as estuaries, tidal pans and salt marshes, emit far less methane into the atmosphere (1.3 g CH_4_ m^−2^ yr^−1^) than freshwater wetlands (7.1 g CH_4_ m^− 2^ yr^−1^) (Bridgham et al., 2013; Bridgham et al., 2006). The lower methane emission is due to the presence of sulfate which enters coastal wetlands via marine seawater inflow and mixes with freshwater inflows creating distinct salinity gradients in the overlying water (Reddy et al., 2022). Salinity in the sediment of coastal wetlands is mostly driven by daily tidal influence (del Pilar Alvarez et al., 2015; Gardner, 2007; Li et al., 2023; Moffett et al., 2010; Reddy et al., 2022) but also wetland geomorphology, sediment characteristics, climate conditions (Montalto et al., 2006), evapotranspiration, bioturbation (Cao et al., 2012; Carol et al., 2011; Li et al., 2023), and anthropogenic alterations (Carol et al., 2012). The sulfate that enters wetlands’ sediment porewater, supports microbial sulfate reduction and thereby suppresses competitive methanogenesis pathways (hydrogenotrophic and acetoclastic) (Jørgensen, 2000; Kristjansson et al., 1982; Lovley and Klug, 1986; Oremland et al., 1982; Oremland and Polcin, 1982; Winfrey and Ward, 1983). As a result, methane tends to buildup in deeper anoxic sediment below the sulfate penetration depth, and thus below the sulfate reduction activity. In the lower portion of the sulfate reduction zone, where the concentration of sulfate is decreasing and overlaps with increasing methane concentration is a sediment zone known as the sulfate methane transition zone (SMTZ), where anaerobic oxidation of methane (AOM) occurs (Hinrichs and Boetius, 2002; Knittel and Boetius, 2009; Reeburgh, 2007). In the SMTZ, AOM oxidizes methane with sulfate as the electron acceptor and has been shown to consume up to 96% of methane before reaching the water column within coastal wetland sediment (La et al., 2022). The process is typically mediated by a consortium of anaerobic methane-oxidizing archaea (ANME) and sulfate-reducing bacteria (Boetius et al., 2000; Hinrichs and Boetius, 2002; Knittel and Boetius, 2009; Michaelis et al., 2002; Orphan et al., 2001; Reeburgh, 2007).

Although sulfate reduction outcompetes methanogenesis for hydrogen and acetate in the sulfate reduction zone above the SMTZ, methylated substrates such as methylsulfides, methanol and methylamines, are known to be non-competitive for methanogenesis of the methylotrophic pathways (Krause et al., 2023; Krause and Treude, 2021; Lovley and Klug, 1986; Maltby et al., 2016; Oremland and Taylor, 1978; Zhuang et al., 2016; Zhuang et al., 2018). Thus, methylotrophic methanogenesis activity has been shown to occur within the sulfate-reducing zone in various aquatic environments, including coastal wetlands (Krause and Treude, 2021; Oremland et al., 1982; Oremland and Polcin, 1982). However, despite the methylotrophic methanogenesis activity, methane concentrations are by several orders of magnitude lower above the SMTZ compared to deeper sediments where sulfate is depleted (Barnes and Goldberg, 1976; Beulig et al., 2018; Krause et al., 2023; Krause and Treude, 2021; Wehrmann et al., 2011). This low level of methane is controlled by concurrent methylotrophic methanogenesis and AOM activity which is now referred as the cryptic methane cycle, and has been detected in marine and coastal wetland sediment (Krause et al., 2023; Krause and Treude, 2021; Xiao et al., 2018; Xiao et al., 2017).

Important questions remain about how methane is cycled in coastal wetlands across spatial gradients and electron acceptor availability. Moreover, the microbial communities, that may be involved directly or indirectly with the cryptic methane cycle have not been identified. Coastal wetlands are ideal geographical features to study these questions because of their unique hydrology where freshwater and marine sources interact creating natural gradients between salinity and terrestrial input, which potentially affects the availability of important electron acceptors (e.g., sulfate, iron (III), nitrate) and the related microbial communities that drive cryptic methane cycling. Therefore, it is crucial to investigate the biogeochemical trends, the metabolic activity of methanogenesis, AOM and sulfate reduction, as well as characterize the microbial communities along a natural salinity gradient of coastal wetlands.

The primary objective is to study cryptic methane cycling along a spatial transect following a natural salinity gradient in a California coastal wetland, the Carpinteria Salt Marsh Reserve (CSMR). Thereby, we aim to elucidate the availability of electron acceptors that drive AOM, and to characterize the microbial communities that are responsible for cryptic methane cycling. In the following study we will show concurrent activities of methanogenesis and AOM, along with potential microbial communities involved, within sulfate-rich sediment across the CSMR land-ocean transect, which strongly suggest active cryptic methane cycling. In addition, our data point to the existence of non-methanogenic anaerobic methylotrophic metabolism alongside the cryptic methane cycle.

## 2. Materials and Methods

### 2.1. Study area and field study

The field site focused on this study is the Carpinteria Salt Marsh Reserve (CSMR), which is located about 15 km east of Santa Barbara, California USA and is part of the University of California Natural Reserve system. Within the CSMR, three freshwater streams flow from the North to the South and open into the Pacific Ocean (Fig. 1). Seawater does infiltrate the CSMR through daily tidal fluxes and mixes with freshwater that enters the CSMR from the North. The mixing results in a natural salinity gradient in the surface waters during low tide. In summer of 2019, sediment samples for this study were collected from a total of four stations (three inside connected creeks and one inside an isolated pool) within the CSMR along a land-ocean, i.e., North-South, transect featuring differences in salinity in the overlying water. Stations were picked based on accessibility and the salinity of the overlying water during low tide, measured in the field with a hand-held refractometer. The stations include brackish low (BL, 7 PSU), brackish high (BH, 15 PSU), marine (M, 35 PSU), and hypersaline (HP, 139 PSU) conditions. The hypersaline conditions are linked to an isolated, evaporative pool, which has been studied previously (Krause and Treude, 2021; Liu, 2024).

**Figure 1.**
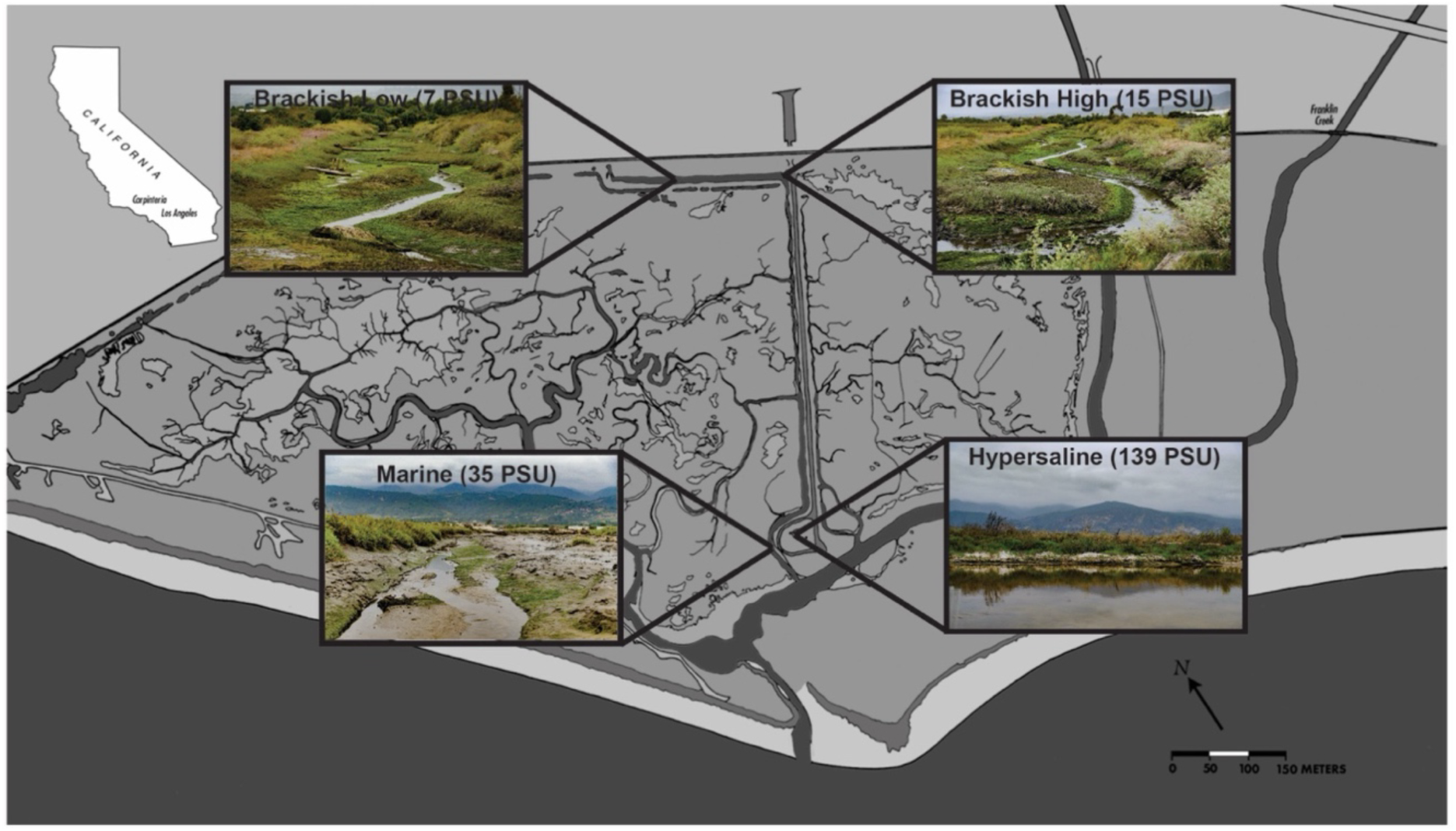
Map of the Carpinteria Salt Marsh Reserve and with sampling locations along the natural salinity gradient. The Brackish Low, Brackish High and Marine stations are connected via creeks; the Hypersaline station is an evaporative isolated pool. PSU = practical salinity units.

The top 15 to 20 cm of sediment at each station was collected in large (10 cm i.d.) and small (2.6 cm i.d.) polycarbonate push cores. Push cores were carefully inserted into sediment by hand including approximately 5 cm overlying water. Large push cores were inserted approximately 20 cm from each other, while small push cores were inserted approximately 15 cm from each other to provide sufficient space for push core excavation and extraction. Sediment surrounding the push cores was carefully removed to place a metal plate under the bottom to safely extract the push cores. Any air headspace within the push cores were filled bubble-free with overlying-water from the station location along the creek or hypersaline pool and sealed with rubber stoppers and electrical tape. Sediment push cores were transported to the home laboratory on the same day, stored in the dark at room temperature and processed within 1 d to 1 week of collection, depending on the analysis type (see 2.2 – 2.7).

### 2.2. Porewater geochemistry and solid phase analysis

One day after collection, one large push core from each station was selected for porewater geochemistry analysis. At all stations the top layer of the sediment was sliced at 1.5 cm followed by 1 cm increments due to natural slopes found at the sediment surfaces during the time of sampling. All sediment was sliced under a constant flow of argon gas to minimize oxidation of oxygen-sensitive substrates. The sediment sections were transferred to pre-argon flushed 50 mL centrifuge vials and centrifuged at 4300 *g* for 20 mins. Immediately after centrifugation, the separated porewater was analyzed spectrographically for dissolved sulfide according to (Cline, 1969) and iron (II) according to (Grasshoff et al., 1999) using a Shimadzu UV-Spectrophotometer (UV-1800) equipped with a sipper unit. The remaining porewater was frozen (-30 °C) and later measured for dissolved porewater sulfate and chloride concentrations. Porewater sulfate and chloride was determined using an ion chromatograph (Metrohm 761) (Dale et al., 2015). Analytical precision of these measurements was <1% based on repeated analysis of IAPSO seawater standards. Absolute detection limit of sulfate was 1 μM, which corresponds to 30 μM in the undiluted sample. Porewater salinity at each station was calculated from chlorinity using Knudson’s equation (Salinity = 1.805 * Chlorinity) assuming that the major ionic ratios in the porewater and in seawater are similar (Knudsen, 1901). One mL of porewater was subsampled for the determination of methylamine concentrations and other methanogenic substrates (see section 2.4).

For methane concentrations, porosity/density, solid-phase carbon/nitrogen, and molecular analysis, a separate large push core from each station, the top 1.5 cm was sliced followed by 1 cm increments because of natural slopes at the sediment surface found at the time of sampling. For methane concentrations, 2 mL of sediment at each interval was subsampled using a 3 mL cut-off plastic syringe and transferred to a 12 mL glass serum vial filled with 5 mL of 5% NaOH and sealed with grey butyl rubber stoppers. Headspace methane concentrations were later determined using a Shimadzu gas chromatography (GC-2014) equipped with a packed Haysep D and flame ionizer detector. The column was heated to 80 °C and ultra-high pure helium was used as the carrier gas, set to 12 mL per minute. A methane standard (Scotty Analyzed Gases) was used to calibrate for methane concentrations with a ± 5% precision.

For porosity and density, 8 mL of sediment was collected from each 1 cm layer using a 10 mL plastic cut off syringe, transferred to pre-weighed plastic 10 mL vials (Wheaton). The wet samples were then weighed and then stored at 4 °C. The samples were later dried at 75 °C for 72 hrs and then reweighed. Sediment porosity was determined by subtracting the dry sediment weight from the wet sediment weight and dividing by the total volume. Sediment density was determined by dividing the wet weight by the total volume of the sample.

Analyses for sediment total organic carbon (TOC) and total organic nitrogen (TON) were modified from (Harris et al., 2001). Briefly, samples were dried up to 48 hours at 50°C until the dry weight was stable and then treated with direct addition of 1 mL of 6N HCl to dissolve carbonate minerals. These samples were then washed in triplicate with 1-mL of ultrapure water or until a neutral pH was re-established. Samples were centrifuged at 4255 X *g* for 20 minutes, the supernatant was decanted, and vials were re-dried at 50°C. A subsample (approx. 10-15 mg) was then packed into individual 8×5 mm pressed tin capsules and sent to the University of California Davis Stable Isotope Facility for analysis using Elemental Analyzer – Isotope Ratio Mass Spectrometry. TOC and TON were calculated based on the sample peak area corrected against a reference material (alfalfa flour).

For molecular analysis, 2 sets of 3 mL of sediments were collected using a 3 mL plastic cut-off syringe into 3 mL plastic cryo vials and immediately stored at -80°C for further analysis (see section 2.9 for details).

### 2.3. *In-vitro* net methanogenesis

One week after sample collection, one large push core from each station was selected to study in-vitro methanogenesis in the natural sediments. Between 2 and 3 sediment intervals at each station were selected and subsampled for this analysis based on porewater geochemistry and solid-phase analysis (see details in Fig. 3).

Sediment subsampling was performed similarly to section 2.2. Each push core was sliced at the designated sediment layers under a constant flow of argon to minimize oxygen poisoning of anaerobic microbial communities within the sediment. Using a 3 mL plastic cut-off syringe, 10 mL sediment from each interval was transferred into triplicate sterilized, argon-flushed 60 mL glass serum vials. Vials were sealed with blue butyl rubber stoppers (Bellco Glass Inc, 20 mm diameter) and crimped with aluminum crimps. The headspace of each vial was flushed with argon for one minute to remove oxygen. The vials were then incubated in the dark, at room temperature and monitored for 22 days. Methane concentrations in the headspace were tracked using a gas chromatograph (see section 2.2).

### 2.4. Metabolomic analysis

Sediment porewater concentrations of methanogenic substrates (methylamine, methanol, and acetate), were obtained from a selection of depth intervals at each station (see specifics in section 3.1.4.) by syringe-filtering (0.2 µm) 1 mL porewater into pre-combusted (350 °C for 3 hrs) amber glass vials (1.8 mL). Samples were then closed with screw caps equipped with a PTFE septa and frozen at -80 °C until analyses. Samples were analyzed at the Pacific Northwest National Laboratory, Environment and Molecular Sciences Division for metabolomic analysis using proton nuclear magnetic resonance (NMR). For details on sample preparation and analysis see section 2.4 in Krause et al. (2023).

### 2.5. Sulfate reduction (^35^S-Sulfate)

Within the one day of collection, one small sediment whole round push core from each station was used to determine sulfate-reduction rates (SRR) at the home laboratory. Radioactive, carrier-free ^35^S-sulfate (^35^S-SO_4_^2-^; dissolved in MilliQ water, injection volume 10 µL, activity 260 KBq, specific activity 1.59 TBq mg^−1^) was injected into the whole-round cores at 1 cm intervals and incubated at room temperature and in the dark following (Jørgensen, 1978). The incubation was stopped after ∼24 hours. Sediment samples (including controls) were transferred, preserved, and stored according to Krause and Treude (2021). Samples were analyzed using the cold-chromium distillation method and the results from the analysis were used to calculate the sulfate reduction rates according to (Kallmeyer et al., 2004).

### 2.6. Methanogenesis and AOM from mono-methylamine

The present study aimed to follow the methane production by methanogenesis from mono-methylamine (hereafter abbreviated MG-MMA) and the subsequent oxidation of the methane to dissolved inorganic carbon (DIC) by anaerobic oxidation of methane (hereafter abbreviated AOM-MMA) (i.e., cryptic methane cycling) in salt marsh sediments across a salinity gradient. To find evidence of concurrent MG-MMA and AOM-MMA, one small whole round core from each station was injected with radiolabeled ^14^C-mono-methylamine (^14^C-MMA) (^14^C-mono-methylamine dissolved in 1 mL water, injection volume 10 µL, activity 220 KBq, specific activity 1.85-2.22 GBq mmol^−1^) at 1-cm intervals according to (Krause and Treude, 2021) and stored at room temperature and in the dark for 24 hrs. Incubations were terminated by slicing the sediment at 1-cm intervals into 50 mL wide-mouth glass crimp vials filled with 20 mL of 5% NaOH. After transfer of the sample, vials were immediately sealed with a red butyl stopper and crimped with an aluminum crimp. Control samples were prepared by sectioning the top 5 cm of a separate whole round core from each station in 1-cm intervals into 50 mL wide mouth vials filled with 20 mL of 5% NaOH prior to radiotracer addition. Vials were shaken thoroughly for 1 min to ensure complete biological inactivity and stored upside down at room temperature till further processing. The residual ^14^C-MMA in the liquid, the ^14^C-CH_4_ in the headspace of the sample vials produced by MG-MMA, and the ^14^C-TIC in the sediments as a result of AOM-MMA samples were determined by the analysis according to (Krause and Treude, 2021).

To account for the ^14^C-MMA binding to mineral surfaces (Wang and Lee, 1993, 1994; Xiao et al., 2022), we determined the recovery factor (RF) for the sediment from stations BL, BH and M following the procedure of Krause and Treude (2021). For the HP station, the RF factor determined by Krause and Treude (2021) was applied.

Estimates of metabolic rates of MG-MMA and AOM-MMA were calculated from the results of the ^14^C-MMA incubations. Natural concentrations of mono-methylamine in the sediment porewater were detectable (> 3 µM) but were below the quantification limit (10 µM) (See section 3.1.4). To enable rate calculations for MG-MMA (Eq. 1), we assumed an MMA concentration of 3 µM for all samples, i.e., the detection limits of the NMR analysis.

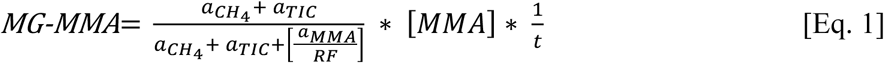

where *MG-MMA* is the rate of methanogenesis from MMA (nmol cm^−3^ d^−1^); *a_CH4_* is the radioactive methane produced from methanogenesis (CPM); *a_TIC_* is the ^14^C-TIC produced from the oxidation of methane (CPM); *a*_MMA_ the residual ^14^C-MMA (CPM); RF is the recovery factor; *[MMA]* is the assumed MMA porewater concentrations (nmol cm^−3^); *t* is the incubation time (d). ^14^C-CH_4_ and ^14^C-TIC sample activity was corrected by respective abiotic activity determined in dead controls.

Results from the ^14^C-MMA incubations were also used to estimate the AOM-MMA rates according to Eq. 2,

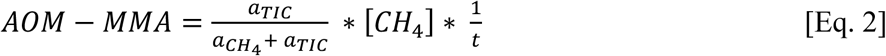

where *AOM-MMA* is the AOM rate based on methane produced from MMA (nmol cm^−3^d^−1^); *a_TIC_* is the produced ^14^C-TIC (CPM); *a*_CH4_ is the residual radioactive methane (CPM); *[CH_4_]* is the sediment methane concentration (nmol cm^−3^); *t* is the incubation time (d). ^14^C-TIC activity was corrected by abiotic activity determined by replicate dead controls.

### 2.7. AOM from ^14^C-methane

AOM rates from ^14^C-CH_4_ (AOM-CH_4_) were determined by injecting radiolabeled ^14^C-CH_4_ (^14^C-CH_4_ dissolved in anoxic MilliQ, injection volume 10 µL, activity 5 KBq, Specific activity 1.85−2.22 GBq mmol^−1^) directly into a separate small whole round core from each station at 1-cm intervals, similar to sections 2.5 and 2.6. Incubations were stopped after ∼24 hours and stored at room temperature until further processing, similar to section 2.6. Sediments were then analyzed in the laboratory using oven combustion (Treude et al., 2005) and acidification/shaking (Joye et al., 2004). The radioactivity captured after the headspace combustion and acidification and shaking analysis were determined by liquid scintillation counting. AOM-CH_4_ rates were calculated according to Eq. 2.

### 2.8. Metabolic rate constants for MG-MMA, AOM-MMA and AOM-CH_4_

Experimental data determined by sect. 2.6 and 2.7 were used to calculate metabolic rate constants (*k)* to compare relative turnover of MMA and CH_4_. We define the rate constants as the metabolic products divided by the sum of the metabolic reactants and products, divided by the incubation time (Krause et al., 2023).

### 2.9. Molecular analysis

#### 2.9.1. DNA extraction from sediment

DNA was extracted from approximately 25-30 mg of sediment collected from all sediment intervals in the top ∼20 cm of the solid phase push cores from all stations (see above section 2.2) using the PowerSoil DNA Extraction Kit (Qiagen) according to the manufacturer’s instructions with the following modifications. Bead tubes were beadbeated for 45 sec at 5.5 m/sec. Samples were eluted first with 50 µL of elution buffer (Solution C6) followed by and additional 25µl of elution buffer for a total of 75uL elution volume.

#### 2.9.2. 16S rRNA gene sequencing (Illumina MiSeq)

The V4-V5 region of the 16S rRNA gene was amplified using archaeal/bacterial primers with Illumina (San Diego, CA, USA) adapters on 5’ end (515F 5’-TCGTCGGCAGCGTCAGATGTGTATAAGAGACAG-GTGYCAGCMGCCGCGGTAA-3’and 926R 5’-GTCTCGTGGGCTCGGAGATGTGTATAAGAGACAG-CCGYCAATTYMTTTRAGTTT-3’). PCR reaction mix was set up in duplicate for each sample with Q5 Hot Start High-Fidelity 2x Master Mix (New England Biolabs, Ipswich, MA, USA) in a 15 μL reaction volume according to manufacturer’s directions with annealing conditions of 54°C for 30 cycles. Duplicate PCR samples were then pooled and barcoded with Illumina Nextera XT index 2 primers that include unique 8-bp barcodes (P5 5’-AATGATACGGCGACCACCGAGATCTACAC-XXXXXXXX-TCGTCGGCAGCGTC-3’ and P7 5’-CAAGCAGAAGACGGCATACGAGAT-XXXXXXXX-GTCTCGTGGGCTCGG-3’). Amplification with barcoded primers used Q5 Hot Start PCR mixture but used 2.5 μL of product in 25 μL of total reaction volume, annealed at 66°C, and cycled only 10 times. Products were purified using Millipore-Sigma (St. Louis, MO, USA) MultiScreen Plate MSNU03010 with vacuum manifold and quantified using ThermoFisher Scientific (Waltham, MA, USA) QuantIT PicoGreen dsDNA Assay Kit P11496 on the BioRad CFX96 Touch Real-Time PCR Detection System. Barcoded samples were combined in equimolar amounts into single tube and purified with Qiagen PCR Purification Kit 28104 before submission to Laragen (Culver City, CA) for 250 bp paired end sequencing on Illumina’s MiSeq platform the addition of 15-20% PhiX.

#### 2.9.3 Analysis of microbial community distribution and 16S rRNA gene sequence data

Sequence data was processed in DADA2 version 1.18 (Callahan et al., 2016). Adapters were removed using cutadept (Martin, 2011). Raw sequences were trimmed to 260bp for forward reads, and 180 bp for reverse reads based on quality of reads. Reads shorter than 260/180 were removed. Error rate was calculated using DADA2’s algorithm. Reads were denoised and merged into Amplicon Sequencing Variants (ASV), requiring a 12bp overlap, and chimeras removed. Taxonomic identification for each representative sequence was assigned with the Silva-138 database (Quast et al., 2012) at 100% identity. The SILVA database had been appended with 1,197 in-house high-quality, methane seep-derived bacterial and archaeal clones. The modified SILVA database is available from the authors upon request.

To visualize the differences between each horizon at each station, a non-metric multidimensional scaling (NMDS) plot was generated. We used Bray-Curtis dissimilarity matrix to calculate the beta diversity of each sample. Each sediment horizon was plotted as an individual dot. The effect of various measured sediment geochemical parameters and calculated metabolic rate measurements for each horizon was calculated with the envfit function (vegan package, permutations = 999, (Oksanen et al., 2022) and plotted as arrows. The arrow vector length corresponds to the significance of its correlation with the NMDS. The length of the arrow corresponds to the significance (all p < 0.05)

## 3. Results

### 3.1. Geochemical trends across the salinity transect

#### 3.1.1 Total organic carbon and nitrogen

Fig. 2 B, H, N, and T show the total organic carbon (TOC) and total organic nitrogen (TON) (wt %) and the C/N (based on the ratio of TOC/TON) ratio in the sediment across the sampled salinity gradient. At stations BL and BH, the TOC profiles varied between 1.4% - 4.4% with the highest TOC near the sediment-water interface (Fig. 2B and H). Note at BH, below 11.5 cm TOC decreased < 0.5%. At the M station, TOC was lower than at stations BL and BH, but generally increased with increasing sediment depth, reaching 3.9% at 14.5-15.5 cm (Fig. 2N). The HP had the highest TOC in the top 1.5 cm (max 5.4%). Below 1.5 cm, TOC sharply decreased to <0.5% (Fig. 2T).

**Figure 2.**
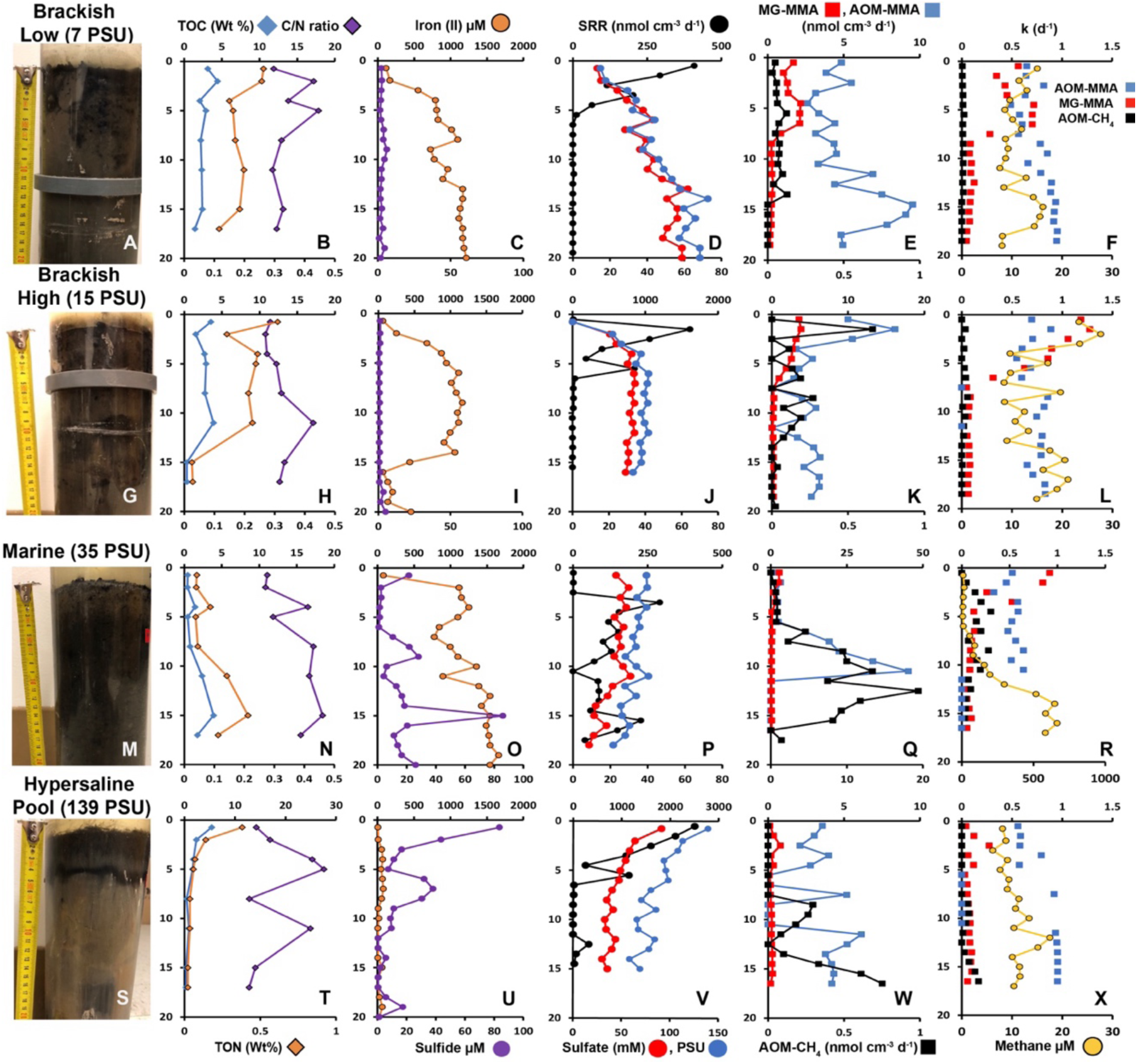
Depth profiles of (bio)geochemical parameters determined in sediment cores collected from the four different stations in the Carpinteria Salt Marsh Reserve. Left Panel: Photos of the sediment cores collected (with scale) from the Brackish Low (**A**), Brackish High (**G**), Marine (**M**), and Hypersaline (**S**) stations. **B**, **H**, **N**, and **T**: TOC, TON and C/N ratio. **C**, **I**, **O**, and **U**: Porewater sulfide and iron (II). **D**, **J**, **P**, and **V**: ex-situ sulfate reduction rates (SRR), porewater sulfate, and porewater salinity. **E**, **K**, **Q**, and **W**: AOM rates derived from ^14^C-mono-methylamine (AOM-MMA) incubations, methanogenesis rates from ^14^C-mono-methylamine (MG-MMA) incubations, and AOM directly derived from ^14^C-methane (AOM-CH4) incubations. **F**, **L**, **R**, and **X**: Metabolic rate constants (k) of AOM-MMA, MG-MMA, AOM-CH4 incubations and sediment methane concentration. Note scale changes on the x-axis.

Generally, TON profiles followed the shape of the TOC profiles at all stations. At BL, TON was between 0.12% and 0.26% (Fig. 2B). At the BH station, TON was between 0.03% and 0.31% (Fig. 2H). At the M station, TON was between 0.04% and 0.21% (Fig. N). At the HP, TON was between 0.02% and 0.38% (Fig. 2T)

At the BL station, the C/N ratio varied between 12 and 18 with the highest C/N within the top 5.5 cm (Fig. 2B). The C/N ratio at the BH station was between 11 and 17 with the highest C/N below 8.5 cm (Fig. 2H). The C/N ratio at the M station increased with sediment depth varying between 11 and 18 (Fig. 2N). The HP station had the highest C/N ratios of all stations, ranging between 13 and 28 with two peaks between 2.5 and 4.5cm and at 10.5-11.5 cm (Fig. 2T).

#### 3.1.2. Salinity and sulfate concentrations

Fig. 2 D, J, P, and V depict the porewater salinity and sulfate concentrations along the sampled salinity gradient. Refractometer measurements of the overlying surface water in the field during low tide showed a natural salinity gradient with fresher and brackish conditions in the northern portion and more saline/hypersaline in the southern portions of the CSMR (Fig. 1). Different to the concentrations found in the overlying surface water, porewater salinity and sulfate increased with sediment depth at both the BL and the BH station. However, max concentrations were lower at BH (47 PSU and 39 mM sulfate) than at BL (73 PSU and 62 mM sulfate) (Fig. 2 D and J). Despite the M station’s proximity to the Pacific Ocean and overlying water salinity resembling marine salinities, porewater salinity and sulfate concentrations at the M station decreased from 40 to 22 PSU and from 30 to 9 mM, respectively, with increasing sediment depth (Fig. 2P). The salinity of the overlying water at the HP exceeded all other stations by one order of magnitude (max 135 PSU), classifying it as hypersaline. The porewater salinity and sulfate concentrations at the HP were the highest at the surface (139 PSU and 91 mM, respectively) but decreased with increasing sediment depth (58 PSU and 30 mM, respectively) (Fig. 2V).

#### 3.1.3. Sediment methane concentrations

At the BL station, methane concentration ranged between 7.6 and 16 µM throughout the core without a clear pattern (Fig. 2F). At the BH station, methane was slightly higher in the top 3.5 cm (max 28 µM). Below 3.5 cm, methane concentrations varied with depth ranging between 7.6 and 21 µM (Fig. 2L). Methane concentrations at the M station were lowest in the top 6.5 cm (7 to 16 µM). Below 6.5 cm, methane steeply increased, reaching 55-665 µM between 6.5 and 19.5 cm, which was the highest methane concentration measured across the salinity gradient (Fig. 2R). The steep increase in methane concentration indicates the shallow end of the sulfate-methane transition zone. However, sulfate concentration was still > 9 mM at these depths, suggesting that the actual transition zone is located much deeper. Methane concentrations at the HP station peaked at 11.5-12.5 cm reaching as high as 17.5 µM, otherwise methane was between 6 µM and 15 µM without a pattern (Fig. 2X).

#### 3.1.4. Substrate availability for methanogenesis

Porewater concentrations of carbon substrates known to support methanogenesis were determined by NMR. Mono-methylamine was detected in the 0-1.5 cm interval at the BL and HP stations but was below quantification limits (10 µM). At all other sediment intervals and stations, mono-methylamine was below the detection limit (< 3 µM). Methanol was detected below quantification (< 10 µM) within the 9.5-10.5 cm intervals at BL and BH stations and the 14.5-15.5 cm intervals at the M and HP stations. Otherwise, methanol was below the detection limits (<3 µM) at all other sediment intervals across all stations. Acetate was present at quantifiable amounts in certain depth intervals at stations BL (9.5 -10.5 cm), BH (0-1.5 cm), and M (14.5-15.5 cm) reaching 61, 45, 72 µM, respectively. Acetate was also detected but below quantification (<10 µM) at stations BH (9.5-10.5 cm) and HP (9.5-10.5 and 14.5-15.5 cm). In all other samples, acetate was below detection (<3 µM).

#### 3.1.5. Total sulfide and iron (II) concentrations

Porewater sulfide (H_2_S, HS^−^, S^2-^) concentrations were consistently low throughout the sediment core at BL and BH, ranging between 0.8 and 6 µM. In contrast, porewater iron (II) concentrations generally increased with increasing sediment depth at BL and BH reaching 1219 µM and 1116 µM, respectively (Fig. 2C and I). Notably iron (II) concentrations at BH decreased from 1116 µM at 10.5 – 11.5 cm to 63 µM at 15.5-16.5 cm before again increasing to 453µM and the bottom of the core (Fig. 2I). Porewater sulfide concentrations were higher at the M station than at the BL and BH stations with peaks at 0-1.5 cm (22 µM), between 6.5 and 10.5 cm (28 µM), and at 15.5-16.5 cm (86 µM). Porewater sulfide concentrations at all other depths were low ranging between 1 to 18 µM (Fig. 2O). Despite having slightly higher sulfide concentrations at the M station, porewater iron (II) concentrations were also high, generally increasing with increasing sediment depth reaching 1661 µM towards the bottom of the sediment core. The HP station showed an inverse relationship between sulfide and iron (II). The highest porewater sulfide concentrations (max 84 µM) were detected at 0-1.5 cm. Below 1.5 cm, sulfide decreased to 7 µM at 4.5-5.5 cm before broadly peaking between 5.5 and 8.8 cm (max 38 µM). Two smaller peaks between 12.5 and 14.5 cm and between 16.5 and 20.5 cm, reached 6 µM and 17 µM, respectively. Iron (II) concentrations varied without trend ranging between 6 µM and 80 µM (Fig. 2U).

### 3.2. Sulfate reduction from ^35^S-sulfate and AOM from ^14^C-CH_4_

Fig. 2D, J, P and V show ex-situ rates of sulfate reduction determined from ^35^S-sulfate incubations across the salinity gradient. At the BL and HP stations sulfate reduction rates were the highest at 0-1 cm, reaching 409 and 2506 nmol cm^−3^ d^−1^, respectively (Fig. 2D and V). Whereas rates at 0-1 cm at the BH and M stations were considerably lower (11 and 1.5 nmol cm^− 3^ d^−1^) (Fig. 2J and P). The rates at the BH and M stations were highest at slightly deeper sediment intervals (at 1-2 cm and at 3-4 cm, respectively), reaching 1615 and 290 nmol cm^−3^ d^−1^. Below the most active sulfate reduction zones at the BL, BH and HP stations, rates decreased with increasing sediment depth to either below detection (i.e., below the control levels) or reaching as low as 4, 0.8, and 10 nmol cm^−3^ d^−1^, respectively. However, at the M station, below 4 cm rates decreased with increasing sediment depth where it was below detection at 9-10cm (Fig. 2P). Below 10 cm rates broadly peaked with the highest rates at 15-16 cm (227 nmol cm^−3^ d^−1^).

Fig. 2E, K, Q, and W show the rates of AOM determined from ^14^C-CH_4_ incubations across the transect (AOM-CH_4_). At the BL station, AOM-CH_4_ rates were low in the top 14 cm ranging between 0.03 and 0.12 nmol cm^−3^ d^−1^ and below detection (i.e., below the control level) below 14 cm (Fig. 2E). At the BH station AOM-CH_4_ rates varied between below detection and 0.66 nmol cm^−3^ d^−1^ at 1-2 cm (Fig. 2K). At the M station, AOM-CH_4_ was either low or absent within the top 6 cm (max 0.9 nmol cm^−3^ d^−1^) but then steeply increased reaching a maximum at 12-13 cm (19.4 nmol cm^−3^ d^−1^) followed by a gradual decreased to 1.3 nmol cm^−3^ d^−1^ at 17-18 cm (Fig. 2Q). At the HP station, AOM-CH_4_ rates in the top 7.5 cm were below detection. This zone of high AOM activity coincided with a steep change in methane concentration (Fig. 2R). Below 7.5 cm, AOM-CH_4_ peaked between 8.5 and 12.5 cm and between 13.5 and 17.5 cm (0.3 and 0.75 nmol cm^−3^ d^−1^, respectively) (Fig. 2W).

### 3.3. Methanogenesis and AOM from ^14^C-MMA

#### 3.3.1. ^14^C-MMA recovery factor (RF)

RF values determined in sediment from BL, BH and M stations (see Sect. 2.6.) were 0.66, 0.62, and 0.57, respectively, and were used to correct MG-MMA rates at each station.

#### 3.3.2. MG-MMA and AOM-MMA

Fig. 2 E, K, Q, and W, show the ex-situ rates of methanogenesis from mono-methylamine (MG-MMA) and AOM (AOM-MMA) determined in ^14^C-MMA incubations and calculated using Equations 1 and 2. Note that MG-MMA rates are only rough estimates assuming a natural MMA concentration of 3 µM, the detection limits of the method, because natural porewater MMA concentrations were either below detection or below quantification (see Sect. 3.1.4.). At BL, MG-MMA varied between 2.1 and 1 nmol cm^−3^ d^−1^ in the top 6.5 cm (Fig. 2E). Between 7 and 19 cm, rates sharply decreased where they remained between 0.14 and 0.36 nmol cm^−3^ d^−1^ without trend. At BH, MG-MMA rates gradually decreased between 0 and 7 cm from max 3.8 to 0.14 nmol cm^−3^ d^−1^ (Fig. 2K). Below 7 cm, rates were mostly stable around 0.2 nmol cm^−3^ d^−1^, except at 11-12 cm, where the rate was below detection, i.e. below the control level. At the M station, MG-MMA rates were overall much lower compared to the BL and BH stations (Fig. 2Q). The highest rate was detected at the sediment surface (0-1 cm, 2.7 nmol cm^−3^ d^−1^) and rates then decreased to <1 nmol cm^−3^ d^−1^ in the remaining core without any trend. At the HP station, MG-MMA rates were the lowest (Fig. 2W). Two peaks were detected between 0 and 3.5 cm and at 4.5-5.5 cm (0.82 and 0.35 nmol cm^−3^ d^−1^, respectively). Below 5.5 cm rates were ranged between 0.08 and 0.3 nmol cm^−3^ d^−1^.

At the BL station AOM-MMA rates, i.e., AOM rates estimated based on the ^14^C-TIC produced during ^14^C-MMA incubations, varied between 2.6 and 5.5 nmol cm^−3^ d^−1^ in the top 11 cm, which are two orders of magnitude higher than AOM rates determined by direct ^14^C-CH_4_ injection (AOM-CH_4_) at this station (Fig. 2E). Below 11 cm, two peaks were detected at 11-12 cm and between 13 and 19 cm, reaching 7 and 9.5 nmol cm^−3^ d^−1^, respectively. At BH, AOM-MMA rates were also higher than AOM-CH_4_, and detected in more sediment intervals, with the highest rates found in the top 3 cm (max 16 nmol cm^−3^ d^−1^) (Fig. 2K). Below 3 cm, rates varied between below detection and 6 nmol cm^−3^ d^−1^. At the M station, AOM-MMA rates were low (max 3.2 nmol cm^−3^ d^−1^) in the top 6 cm, but still an order of magnitude higher than the AOM-CH_4_ (Fig. 2Q). Rates then increased (max 45.2 nmol cm^−3^ d^−1^) between 6 and 11 cm, similar to the AOM-CH_4_ rates. Below 11 cm, the AOM-MMA rates were below detection. At the HP station AOM-MMA rates varied with increasing sediment depth ranging between 2 and 6 nmol cm^−3^ d^−1^ without trend (Fig. 2W). It is notable that AOM-MMA was detected in the top 5 cm while AOM-CH_4_ was not. Conversely, between 5.5 and 7.5 and between 8.5 and 11.5 cm where AOM-CH_4_ was detected, AOM-MMA activity was below the dead control threshold and thus was found to be below detection.

### 3.4. Metabolic rate constants

Metabolic rate constants (k) were plotted to allow for a more direct comparison between the turnover of different ^14^C radiotracer species. At BL, k for AOM-CH_4_ was < 0.1 d^−1^ throughout the sediment core (Fig. 2F), while k for MG-MMA was the highest between 0 and 7 cm (max 0.72 d^−1^). The k for AOM-MMA was high throughout the sediment core and only slightly increased with sediment depth from 0.5 d^−1^ at the surface to 0.9 d^−1^ at the bottom of the sediment core.

The profiles of the three different rate constants looked very similar at BH station when compared to BL. The k for AOM-CH_4_ was constantly low at < 0.1 d^−1^ throughout the sediment core (Fig. 2L). Note at some sediment intervals AOM-CH_4_ rates were not detected and thus rate constants are reported as zero. The k for MG-MMA was high at the surface (1.2 d^−1^ at 0-1cm) and decreased to 0.3 d^−1^ at 6-7 cm. Below 7 cm, k varied slightly between 0.02 and 0.08 d^−1^. The l for AOM-MMA was high throughout the core and varied only slightly with sediment depth between 0.5 d^−1^ and 0.8 d^−1^. Note that also here some ks were zero where AOM-MMA was not detected.

At the M station, k for AOM-CH_4_ was zero at 0-1 cm but increased from 0.05 d^−1^ at 1-2 cm to 0.30 d^−1^ at 4-5 cm. Below 5 cm, k varied between 0.07 d^−1^ and 0.3 d^−1^ (Fig. R). The k of MG-MMA showed similar patterns to BH, i.e., gradually decreasing with increasing sediment depth from 0.92 d^−1^ at 0-1cm to 0.054 d^−1^ 16-17cm. The k of AOM-MMA ranged between 0.33 d^− 1^ and 0.64 d^−1^ in the top 11 cm. Below 11 cm, AOM-MMA rates were not detected thus k was zero at these depths.

At HP, k for AOM-CH_4_ ranged between 0.01 d^−1^ and 0.16 d^−1^ in the 7-11 cm and 13-17 cm sections (Fig. 2X). Otherwise, AOM-CH_4_ rates were not detected thus k is zero for those depths. The k of MG-MMA ranged between 0.04 d^−1^ and 0.27 d^−1^ with a small peak at 2-3 cm. The k of AOM-MMA, ranged between 0.56 d^−1^ and 0.8 d^−1^ in the top 5.5 cm. At 7.5-8.5 cm, and between 11.5 and 17.5 cm k was consistently around 0.95 d^−1^. In between these positive values, k was zero, because no activity was detected.

### 3.5. Methanogenesis batch incubations

Fig. 3 shows results from the in vitro time-series incubation of methanogenesis with natural sediments collected from each station (see details in section 2.3.). Methane in the headspace of vials from all stations and depth intervals did not show any linear increase over the course of 530 hours. At the BL, BH, and HP stations, methane remained very low with small oscillations ranging between 40 and 258 ppmv (Fig. 3A, B, and D).

In the 0-5 cm depth interval of the M station, methane showed a similar pattern as the other stations, ranging from 66 to 115 ppmv (Fig. 3C). However, the 5-14 cm (239 to 356 ppmv) and 14-19 cm (366 to 508 ppmv) interval of the M station showed elevated methane, which peaked at 312 hrs after which methane decreased again.

### 3.6 Microbial diversity and Spatial community patterns

Variation in the microbial community structure from 16S rRNA iTAG analysis (including bacteria and archaea) was observed between the 4 stations, (Fig. 4), with depth affiliated clustering of communities from the BL and BH stations relative to the M and HP stations. NMDS results showed that the communities within each horizon from the BL and BH grouped closer together, indicating similar beta diversity. Interestingly, microbial community structure at the M station showed two distinct groups, split by sediment depth. The microbial community structure at the HP station on the other hand, had the largest spread and was the least similar to the microbial communities from rest of the stations. To determine whether there were correlations between microbial community structure and corresponding geochemical and process rate data, laboratory incubation rates of matching cores as well as environmental factors were fitted onto the ordination using envfit command from the vegan R package (Oksanen et al., 2022). From this analysis, rates of MG-MMA were found to be significantly correlated with microbial communities in the BL and BH stations, while microbial community structure from station M was correlated with AOM-CH_4_ and AOM-MMA rates (P-values = 0.001 and 0.007, respectively), and the HP community was correlated with sulfate reduction (P-value = 0.001).

**Figure 3.**
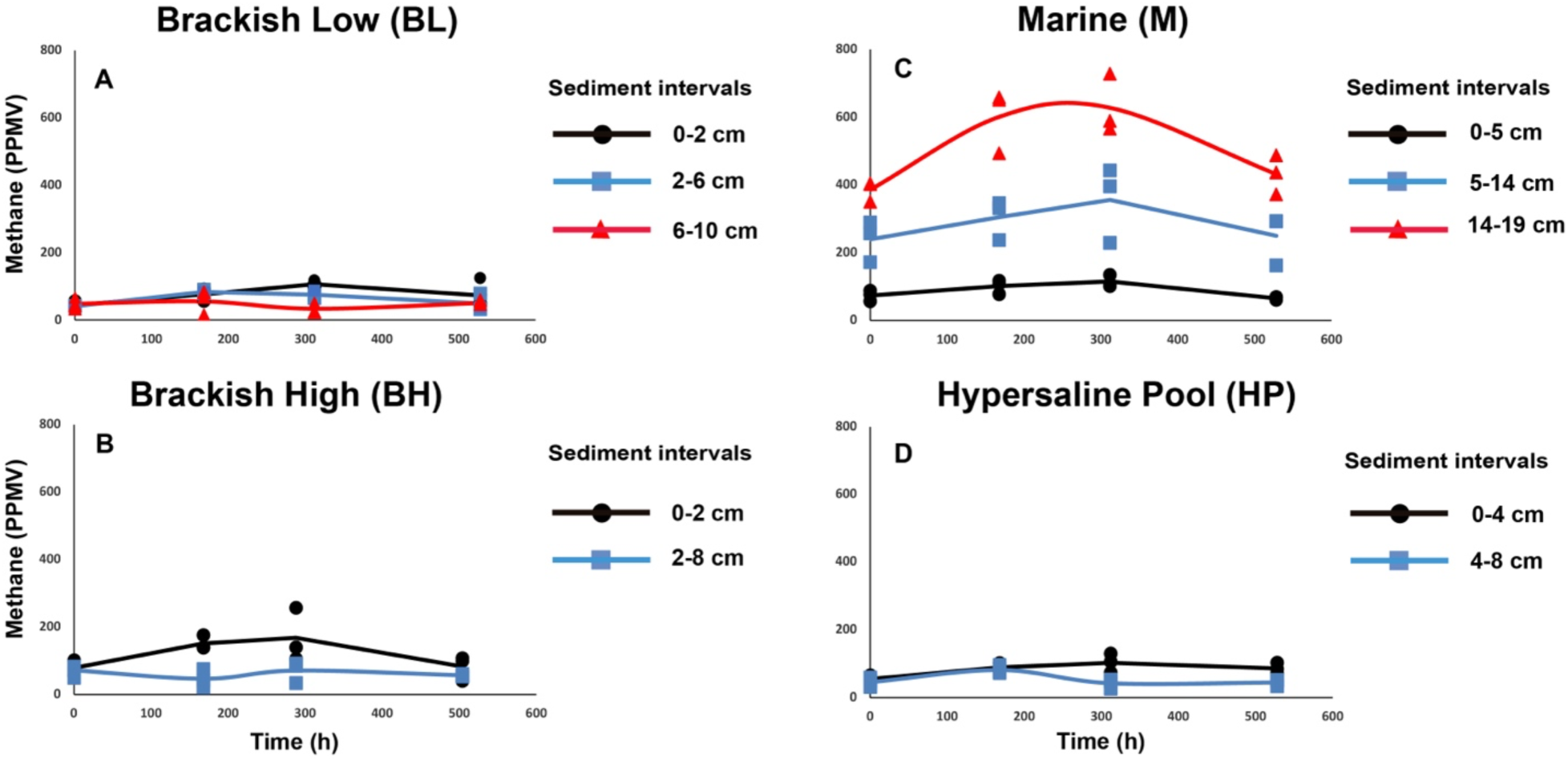
Methane development during methanogenesis batch incubations with sediments from discrete layers over time; (**A**) Brackish Low, (**B**) Brackish High, (**C**) Marine, and (**D**) Hypersaline Pool.

**Figure 4.**
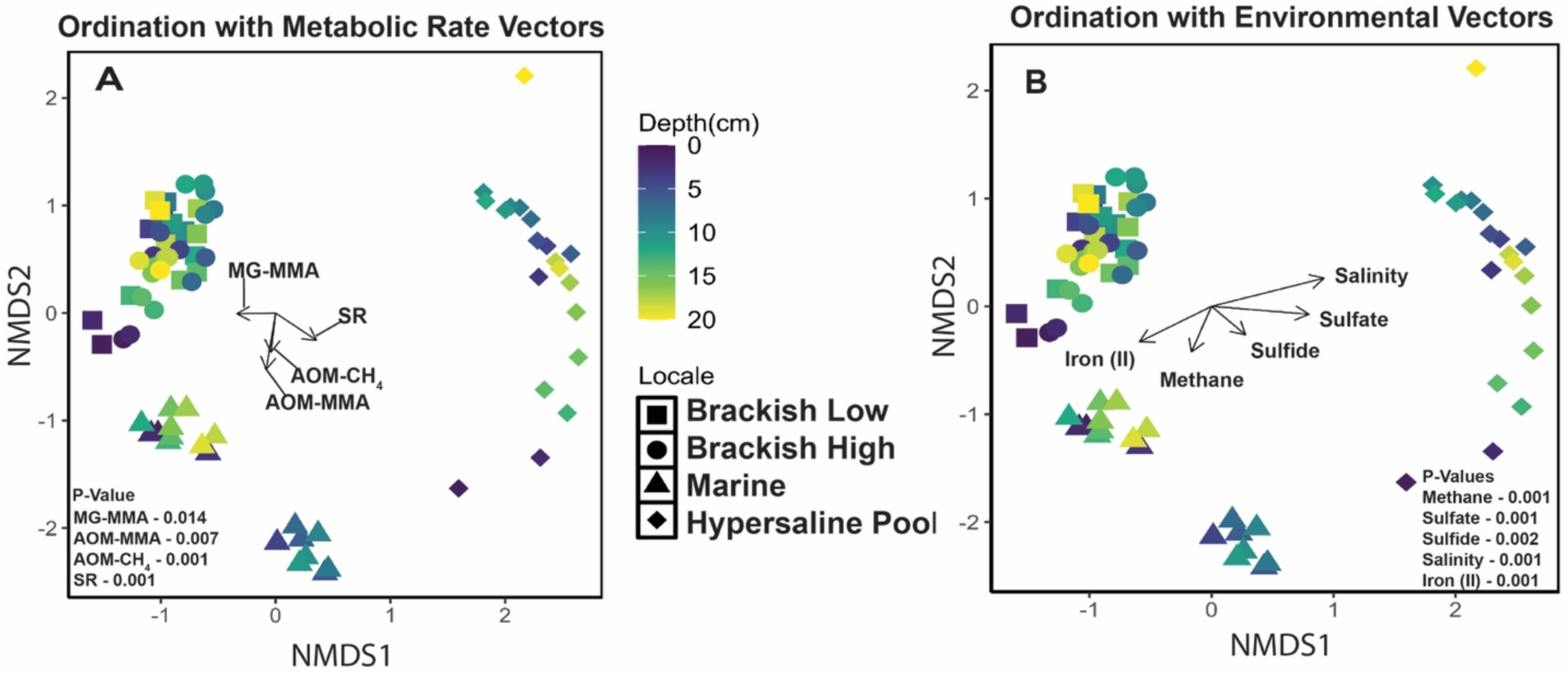
NMDS plots showing microbial community structure of each station relative to each other, sediment depth, and correlations of microbial communities to metabolic rates (**A**) and geochemical parameters (**B**).

Correlations were also observed between community structure and major geochemical parameters, including elevated salinity and sulfate at the HP site (P-value = 0.001), methane concentrations with the microbial community at station M (P-value = 0.001), iron (II) for BL and BH communities (P-value = 0.001), and sulfide for stations M and HP (P-value = 0.002).

### 3.7. 16S rRNA iTAG diversity

16S rRNA iTAG sequencing was used to characterize the archaeal (Fig. 5A-D) and bacterial (Fig. 5 E-H) diversity (above 1% relative abundances), with emphasis on methane related archaea and sulfate-reducing bacteria at each station within the CSMR transect (see sections 3.7.1 and 3.7.2). At the BL and BH stations, the most abundant archaeal amplicon sequence variants (ASVs) were found in sediment intervals between 6.5 and 20.5 cm and between 2.5 and 20.5 cm, respectively (Fig. 5A and B). These ASVs predominantly belonged to archaeal families of Bathyarchaeia (1-3.6% at BL; 1.1 – 8.1% at BH), Lokiarchaeia (1.2-4.8% at BL;1% at BH), Marine Benthic Group D and DHVEG-1 (1.4 – 5.7% at BL; 1.2-4% at BH), and uncultured families within Thermoplasmatota (0.05-1.85% at BL; 0.06-1.25% at BH). At the M station archaeal ASV’s increased with increasing sediment depth and were most abundant in deeper sediment intervals between 11.5 and 19.5 cm (Fig. 5C). In these sediment intervals, archaeal families mostly belonged to Woesearchaeales (1.2-9.8%) and the SCDC AAA0110D5 family of the Nanoarchaeota phylum (1.3-10.4%). At the HP station archaeal ASVs were highest out of all stations and decreased in abundance with increasing sediment depth (Fig. 5D). Here ASV’s belonging to a variety of families of Haloferacaceae were detected up to 33% between 1.5 and 9.5 cm. Below 9.5 cm, one family of Haloferacaceae (Haloferacaceae_1) ASVs were detected at lower relative abundances (1.2-9.4%) but were also accompanied by Nitrosopumilaceae (1.2-10.7%).

**Figure 5.**
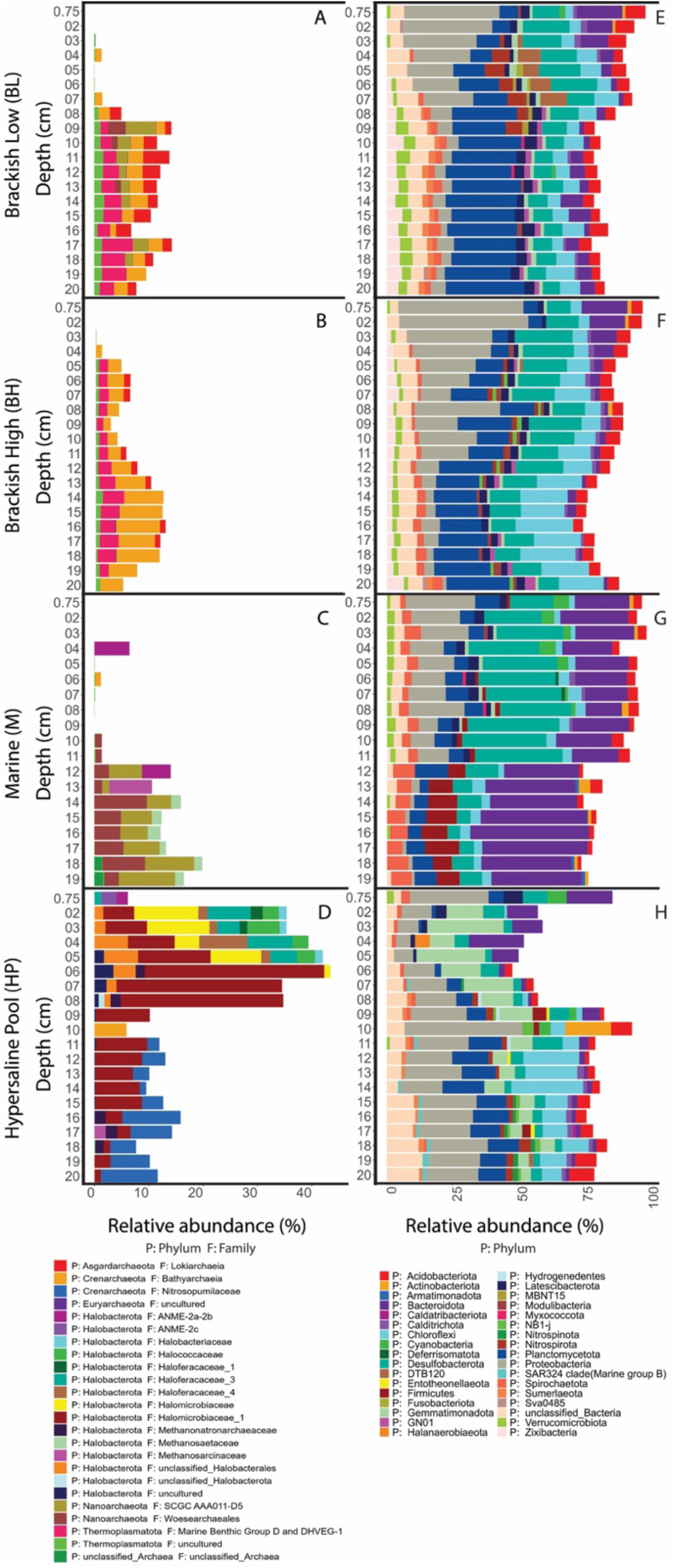
16S rRNA iTAG relative abundances of archaeal phyla above the 0.1% relative abundance threshold (A-D) and of the top 36 bacteria phyla (E-H) detected in the sediment of the CSMR transect. Note the differences in x-axis scale.

Fig. 5 also shows the top 34 bacterial phyla ASVs detected in the sediment across the salinity gradient. Aside from the known bacterial phyla containing bacteria that can perform sulfate/iron reduction, other bacterial phyla seem to be much more dominant in the CSMR sediment across the salinity gradient. Bacterial ASVs belonging to the Pseudomonadales (formerly Proteobacteria) were present throughout all sediment depths from all stations but more prevalent in the top 10.5-11.5 cm at BL, BH, and M stations (35%-26%), and were abundant throughout the sediment at the HP station (5.5%-45%). At the BL and BH stations bacterial ASVs belonging to Planctomycetota are more prevalent in sediment intervals below 7 cm, reaching 29% and 24%, respectively. At the M station, ASVs belonging to Planctomycetota are present throughout the sediment but are lower than at the BL and BH stations, reaching 12%.

#### 3.7.1 Methanogenic and methanotrophic signals

A main goal of this study was to identify methanogenic and methanotrophic archaea that could potentially be contributing to methane cycling across the transect. Fig. 6A-G shows ASVs of the most abundant groups of methane related archaea (above 0.1% relative abundances) recovered from sediment samples across the transect, including known methanogens (*Methanosarcinaceae*, *Methanosaetaceae, Methanoregulaceae, Methanomassiliicoccales*, Methanofastidiosales, and *Methanonatronarchaeacea*) and anaerobic methanotrophs (Candidatus orders Methanocomedens, Methanomarinus, and Methanogastraceae (ANME 2a-c))(Chadwick et al., 2022). The groups of methanogens listed above can be found in anoxic sediments within a wide range of environments such as and marine environments, acidic peat bogs, anaerobic reactors, oil fields, rice paddy soil, a mud volcano, and in a freshwater lake , and hypersaline environments (Oren, 2014; Sorokin et al., 2018). These methanogens produce methane using substrates such as acetate (*Methanosarcinaceae* and *Methanosaetaceae*), methylated substrates (*Methanosarcinaceae*, *Methanomassiliicoccales*, and *Methanonatronarchaeacea*), and/or H_2_/CO_2_ (*Methanosarcinaceae*, *Methanoregulaceae*, *Methanomassiliicoccales*, and *Methanonatronarchaeacea*) (Liu and Whitman, 2008; Oren, 2014; Sorokin et al., 2018). It is worth noting that iron-dependent methanotrophic growth have been observed in groups within *Methanosarcinaceae*, implicating them in iron-dependent AOM (Yan et al., 2023; Yan et al., 2018)

**Figure 6.**
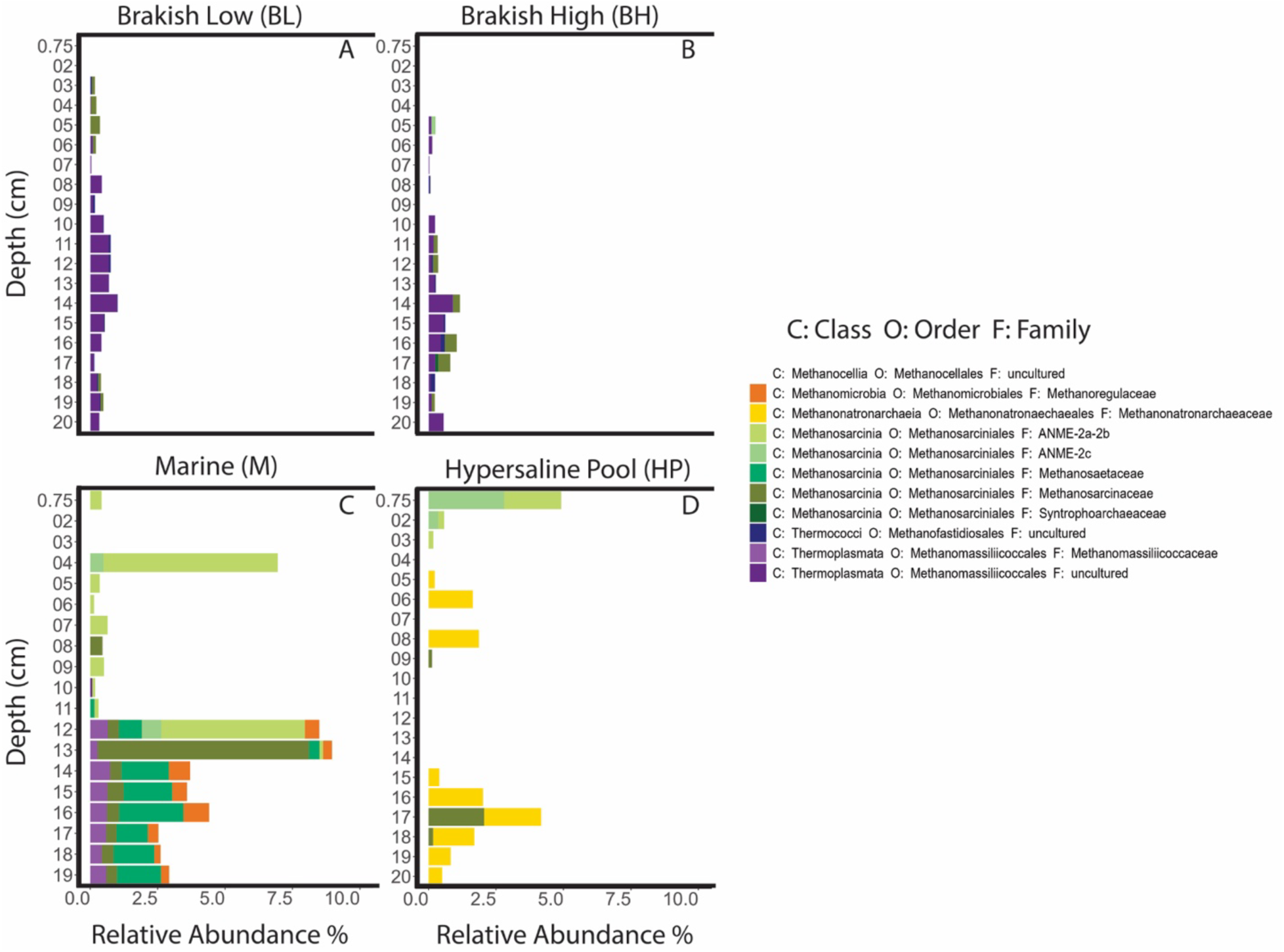
Relative abundances of putative methanogenic and anaerobic methanotrophic archaea (A-D) and the sum (E-H) along the CSMR transect.

The Ca. Methanocomedens and Ca. Methanomarinus (ANME 2a and b, respectively) lineage are commonly found in diverse methane-rich deep-sea and coastal marine environments (Ruff et al., 2016) and salt marsh sediment, mud volcanoes, hydrothermal vents (ANME 2a), and frequently co-exist with members of the Methanogasteraceae (ANME-2c) in methane seeps (Knittel and Boetius, 2009; Knittel et al., 2018). ANME-2c has additionally been reported from a freshwater coastal aquifer (López-Archilla et al., 2007). All of these methanotrophic ANME-2 have been found to partner with diverse syntrophic sulfate-reducing bacteria to couple AOM with sulfate reduction (Knittel and Boetius, 2009; Knittel et al., 2018; Murali et al., 2023; Timmers et al., 2017).

The BL and BH stations had the lowest relative abundance of archaea putatively associated with methane metabolism (Fig. 6A and B), with overall relative abundances <2.5% from the families of Methanosarcinaceae and <1% relative abundance of uncultured *Methanomassiliicoccales*, and uncultured Methanofastidiosales. At BL, *Methanosarcinacea* was found mainly at the top horizons (3-6cm), while at BH, low ASV abundances of *Methanosarcinaceae* were recovered from deeper sediment intervals at 13.5-14.5 cm and between 15.5 and 17.5 cm (between 0.26 and 0.46%) (Fig. 6B). At the BH station ASVs affiliated with one group of anaerobic methanotrophic archaea (ANME-2C) was detected at 5.5-6.5 cm (0.15%). Otherwise, no other groups of ANME were detected in the brackish BL and BH stations, which is consistent with low AOM activity measured at these stations (Fig. 2E and K). At the M station, the relative abundance of ASV’s affiliated with both methanogenic and methanotrophic archaea were more prevalent than at any of the other stations (Fig. 6C), here also corresponding with some of the highest AOM rate measurements (Fig. 2Q). Methanotrophic ANME-2a and b were dominant over methanogenic archaeal lineages, reaching 6.5% and 5.3% at 4-5 cm and 11.5-12.5 cm, respectively compared to 0.45% for *Methanosarcinaceae* at 7.5-8.5 cm and 2.3% for the uncultured *Methanomassiliicoccales* at 9.5-10.5 cm. ANME-2c ASVs were also detected at low relative abundance from the 4.5-5.5 and 12.5-13.5 cm reaching 0.48 and 0.73%, respectively. At 12.5-13.5 cm, ASVs belonging to methanogenic archaeal families of *Methanosarcinaceae*, *Methanosaetaceae*, *Methanomassiliicoccales*, and *Methanoregulaceae*, reached 0.41, 0.85, 0.65, and 0.53%, respectively, alongside with ANME 2a-2b and 2c archaeal ASVs at this depth interval (Fig. 6C). Note at this sediment horizon we detect estimated rates of MG-MMA (0.17 nmol cm^−3^ d^−1^), the highest AOM-CH_4_ (19 nmol cm^−3^ d^−1^) (Fig. 2W), and elevated methane concentrations (297 µM) (Fig. 2X). Below 13.5 cm, methanotrophic archaea were not detected, but a higher percentage of the ASV belonged to *Methanosarcinaceae*, *Methanosaetaceae*, *Methanomassiliicoccales*, *and Methanoregulaceae* reaching 7.8, 2.3, 0.74 and 0.95%, respectively (Fig. 6C).

At the HP station, ASVs belonging to ANME 2a-2b, and 2c were detected in the top 2.5 cm, reaching 2.8 and 2.1%, respectively. This coincides with detectable AOM-MMA activity but not with AOM-CH_4_ activity, which were below detection at these depth intervals (Fig. 2W). Below 2.5 cm, methanotrophic ASVs were not recovered (Fig 6D), but AOM was detectable (Fig. 2W). Methanogenic ASVs belonging to the family *Methanonatronarchaeacea,* a methylotrophic methanogenic group previously described from hypersaline environments (Sorokin et al., 2018),were detected in deeper sediment intervals, where MG-MMA activity was also detected (Fig. 2W), between 4.5 and 8.5 cm and again between 15.5 and 20.5 cm reaching as high as 1.9% and 2.1%, respectively (Fig. 6D). *Methanosarcinaceae* ASV’s were also detected at 17.5 – 18.5 cm.

#### 3.7.2 Distribution of sulfate-reducing bacteria

Figure 7A-D shows the ASVs belonging to known bacterial groups capable of sulfate reduction that were detected in the sediment across the CSMR transect. Members within the Desulfobacterota phylum are the dominant sulfate-reducing organisms that are found in surface sediments in marine and freshwater environments (Diao et al., 2023; Waite et al., 2020). It is worth noting that many members within this phylum (Geobacterales, Geopsychrobacteraceae, Geothermobacteraceae, and Desulfuromondaceaae) are able to utilize oxidized metals such as iron as an electron acceptor for their metabolisms (Holmes et al., 2004; Waite et al., 2020; Young, 2003). Furthermore, it has been shown that orders within Desulfobacterota (Desulfobacterales, Desulfobulbales, Desulfovibrionales) have members that are able to do iron reduction in addition to sulfate reduction (Park et al., 2008; Reyes et al., 2017)

**Figure 7.**
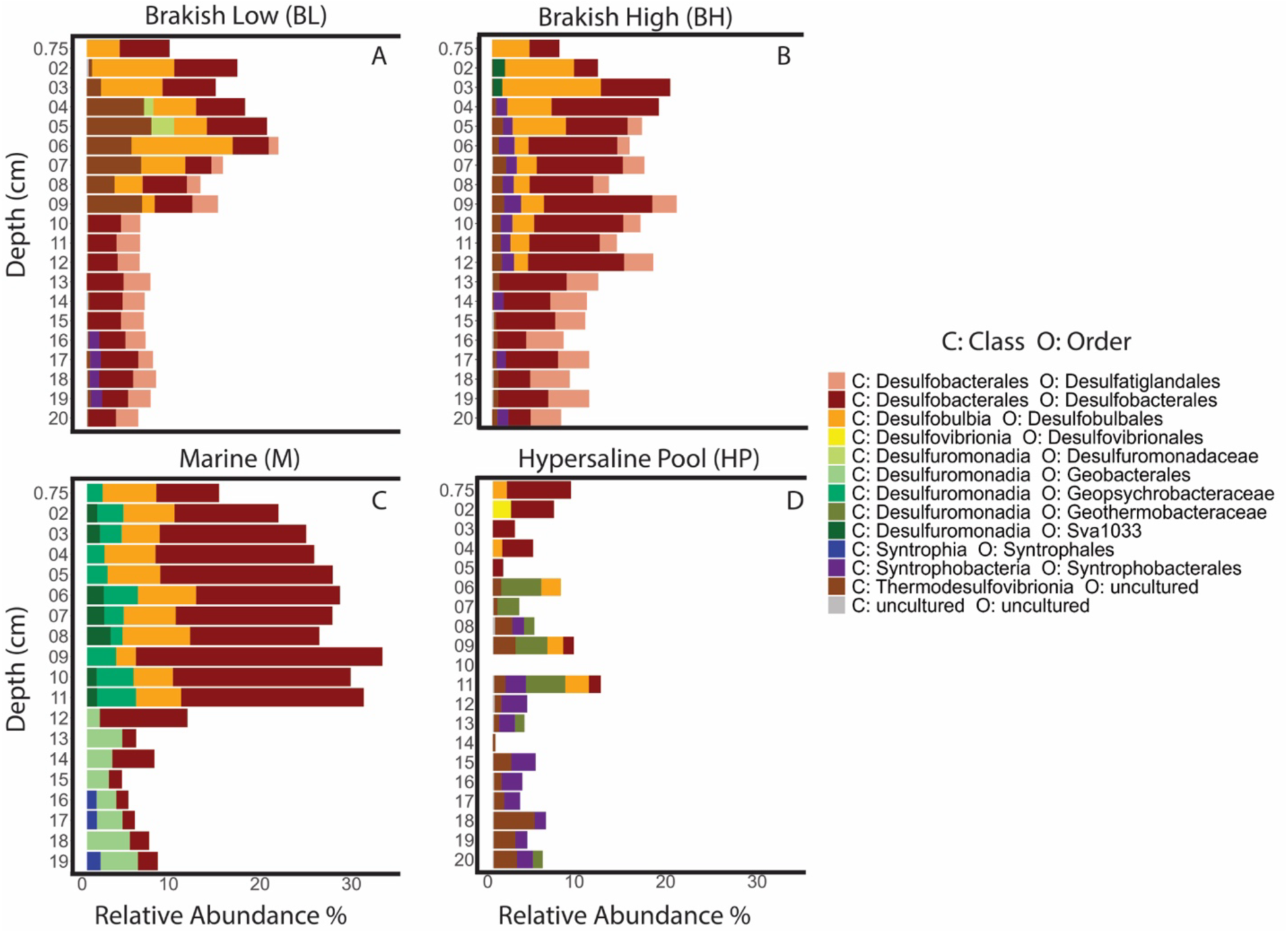
Relative abundances of bacterial classes and orders within the Desulfobacterota phylum.

At the BL station, ASV’s belonging to Desulfobulbales order were detected in the top 9.5 cm (where high sulfate reduction rates and porewater iron (II) were detected (Fig. 2C and D)) with relative abundances ranging between 1.4 and 11 %. Whereas ASVs belonging to Desulfobacterales (part of Desulfobacterota 1) with relative abundances between 2.8-6.9 % were detected throughout the sediment core (Fig. 7A). BL was the only station containing ASV’s belonging to Desulfuromonadaceae between 4.5-6.5 cm ranging between 1.5% and 3.4%. Between 15.5 cm and 19.5 cm ASV’s belonging to Syntrophobacterales were detected at about 1%. At BL between 1.5 cm and 9.5 cm ASV’s belonging to Thermodesulfovibrionia were between 0.4% and 7%. Desulfobulbales and Desulfobacterales ASV trends at the BH station were similar to the BL station, however, Desulfobulbales abundances (ranging between 1.5-10%) extended from the surface into deeper sediment intervals down to 12 cm (Fig. 7B). Desulfobacterales’ ASV’s at the BH station were also higher than at the BL station, ranging between 2.4-11.8%. Stations BL and BH were the only stations with ASV’s belonging to Desulfatiglandales ranging between 1% and 4.5%, which were found between approximately 5.5 cm and 20.5 cm. Between 1.5 cm and 3.5 cm ASV’s belonging to Desulfuromonadia were detected (∼1.5%). ASV’s belonging to Thermodesulfovibrionia were also detected between 3.5 cm and 12.5 cm at low abundances between 0.5% and 1.5%. ASV’s belonging to Syntrophobacterales were also detected at BH between 4.5 and 13.5 cm and at 14.5-15.5 cm, at 17.5-18.5 cm, and 19.5-20.5 cm, ranging between 1% and 1.8%. At the M station, Desulfobacterales abundances were the highest in the top 11 cm, reaching 26.9% at 8.5-9.5 cm (Fig. 7C), and corresponds with high sulfate reduction activity and porewarter iron (II) (Fig. 2I and J). Below 12 cm, Desulfobacterales abundances declined, ranging between 1.3-4.6%. Desulfobulbales was also detected in the top 11 cm but were lower (4.1%-7.4%) compared to the relative abundance of Desulfobacterales. The M station was the only station containing ASV’s belonging to Desulfuromonadia (ranging between 1.3% and 4.3%), Geobacterales (ranging between 1.4% and 4.7%), and Syntrophales (ranging between 1% and 1.4%) in the top 11.5 cm, below 12.5 cm, and below 16.5 cm, respectively. The HP station had the least amount of ASVs belonging to phyla capable of sulfate reduction (Fig. 7D), despite having the highest sulfate reduction activity out of all stations (Fig. 2V). Desulfobacterales were detected between 0 and 5 cm, at 8.5-9.5 cm and at 10.5-11.5 cm ranging between 1.1-7%. Desulfobulbales ASV’s were also detected at 0-1.5 cm, 3.5-4.5 cm, 8.5-9.5 cm and 10.5-11.5 cm with low abundances ranging between 1-2%. One sediment interval at 1.5-2.5 cm had ASV’s belonging to Desulfovibrionales at low relative abundance (2%) (Fig. 7D). The HP station was the only station containing ASV’s belonging to Geothermobacteraceae between 6.5cm and 13.5 cm and at 19.5-20.5 cm, ranging between 1% and 4.3%, where smaller concentrations of porewater iron (II) were detected (Fig. 2U). Between 5.5 cm and 20.5 cm ASV’s belonging to Thermodesulfovibrionia were detected ranging between 0.4% and 4.5%. ASV’s belonging to Syntrophobacterales was also detected in the HP station at 8.5-9.5 cm and between 11.5 and 20.5 cm, ranging between 1.2% and 2.8% (Fig. 7D).

## 4. Discussion

### 4.1 Spatial evidence of cryptic methane cycling

The aim of this study was to find evidence of cryptic methane cycling across the land-ocean transect in the CSMR, by presenting geochemical evidence along with concurrent activity of methanogenesis linked to mono-methylamine (in the following termed ‘methylotrophic methanogenesis’), sulfate reduction, and AOM from radiotracer incubations (^14^C-MMA, ^35^S-SO_4_^− 2^, and ^14^C-CH_4_, respectively). Based on the development of ^14^C-methane in our ^14^C-MMA incubations, methylotrophic methanogenesis appears to be active at all stations and throughout the sampled intervals. Radiotracer experiments further suggested that methylotrophic methanogenesis overlapped with AOM in the 0-14, 1-19, 1-18, and 7-17 cm section at the BL, BH, M, and HP stations, respectively (Fig. 2E, K, Q, and W). Together these findings indicate that methylotrophic methanogenesis is directly fueling AOM at the investigated stations. Note however, that AOM was below the detection limit in some sediment intervals at the BH and M stations and comparably low (< 1 nmol cm^−3^ d^−1^) at the BL and BH stations. Sulfate reduction, a process commonly linked to AOM in marine environments, was found to be active alongside methylotrophic methanogenesis and AOM in the 1-6, 1-16, 0-18 and 13-15 cm sections at the BL, BH, M, and HP stations, respectively (Fig. 2D, E, J, K, P, Q, V, and W). Our data suggest that the activity of sulfate reduction and methylotrophic methanogenesis is driven by the availability of organic matter. For example, higher TOC (3.5-4.4%) and TON (0.26-0.31%) were found within the top 2.5 cm at both BL and BH (Fig. 2B and H). These TOC and TON contents were within the range of TOC and TON found in surface sediment in other salt marshes, such as the North Inlet salt marsh South Carolina, USA (1.6-9.4% for TOC; 0.15-0.6% for TON) (Chen et al., 2016)), the low salt marshes of the Finistere region, France (Goslin et al., 2017), and the salt marshes within the Cádiz Ba, southern Spain (approx. 3% for TOC; approx. 0.3% for TON) (de los Santos et al., 2023).The C/N ratios at the BL and BH stations suggest that the organic matter is likely sourced from a mixture of terrestrial plants (C/N > 20) and marine organic matter (C/N between 5 and 8) (Fig. 2B and H) (Meyers, 1994; Xia et al., 2021). Terrestrial plants were found lined against the embankments and submerged within streams at the BL and BH stations which are likely supplying the organic matter detected (Fig. 1). Previous studies have shown that coastal marsh vegetation is an important source of organic carbon into the sediment (Bartoli et al., 2021; Reddy et al., 2022), which can enhance sulfate reduction (Jackson et al., 2014; Kostka et al., 2002) and methanogenesis (Yuan et al., 2014; 2019).

Accordingly, a lower TOC and TON content was detected at the M station (Fig. 2N), potentially explaining lower or undetectable sulfate reduction and methylotrophic methanogenesis activity within the top 10 cm (Fig. 2P and Q). Field and lab observations of the sediment from the M station indicated a larger sediment grain size (sand) in the top 10 cm compared to finer grained silt and muds found at BL and BH. Previous work have shown that permeable sands are sites for enhanced organic matter consumption (Boudreau et al., 2001) because flow into permeable sandy sediment delivers more particulate organic matter (Huettel et al., 1996; Rusch et al., 2001), and oxygen (Booij et al., 1991; Precht et al., 2004; Ziebis et al., 1996), and quickly remove products of degradation (Huettel et al., 1998; Rocha, 1998). This flushing process likely prevents sediment near the surface from becoming fully anoxic, inhibiting sulfate reduction and methanogenesis at the M station.

Different to Krause and Treude (2021), who found concurrent peaks of methylotrophic methanogenesis and AOM in the top 5 cm at the HP station, overlap of the two processes was detected only in deeper sediment intervals (> 12 cm, Fig. 2W) in the present study. It is worth noting that in the present study sediment from the HP pool was collected approximately one year after and one meter away from the original sampling spot studied in Krause and Treude (2021), suggesting that cryptic methane cycling activity is subject to natural temporal and spatial variation.

Methane concentrations detected within sulfate-rich sediment is likely sourced from methanogenesis via methylotrophic pathways (King et al., 1983; Maltby et al., 2016; Oremland and Taylor, 1978; Reeburgh, 2007), which is further supported by the ^14^C-methane produced during the ^14^C-MMA incubations (Fig. 2 E, K, Q, W). Methane concentrations in sediment cores from the BL, BH, and HP station were about an order of magnitude lower (<28 µM) than what has been previously reported from the top 20 cm of sediment at other coastal wetlands, such as within the Arne Peninsular salt marsh, UK (up to 100 µM) (Parkes et al., 2012), the Queen’s Creek Tidal Marsh, VA, USA (up to 450 µM, in July) (Bartlett et al., 1987), the Chongming Island of the Yangtze Estuary, East Asia (up to 156 µM) (Li et al., 2021), and Dover Bluff salt marsh, GA, USA (0.3-1.2 mM) (Segarra et al., 2013). Instead, methane profiles at the BL, BH and HP stations, were consistently low between 3 and 28 µM (Fig. 2 F, L, and X). These low methane concentrations could represent the equilibrium between methylotrophic methanogenesis and AOM as part of the cryptic methane cycle in sulfate-reducing sediment (Krause et al., 2023). The observed methane profiles are opposite the classical diffusion-controlled trend where residual methane linearly declines between the SMTZ and the sediment-water interface, which is often seen in salt marshes, tidal marshes and estuaries (Bartlett et al., 1987; Parkes et al., 2012; Segarra et al., 2013) and marine environments (Bernard, 1979; Iversen and Jørgensen, 1993; Reeburgh, 2007; Tilbrook and Karl, 1995) including the M station from the present study. Methane profiles at the BL, BH, M (top 5 cm), and HP stations were more similar to organic-rich surface sediment in the oxygen-deficient Santa Barbara Basin, where the simultaneous production and consumption of methane was also detected (Krause et al., 2023). Note the methane profile in deeper sediment intervals (> 5 cm) at the M station resembled classic methane profiles in deeper anoxic sediments found in marine sediment (Bernard, 1979; Reeburgh, 2007; Tilbrook and Karl, 1995). Additional support for the presence of cryptic methane cycling comes from the methanogenesis batch incubations (Fig. 3A, B, C and D), which show no linear buildup of methane in the headspace over ∼530 hrs, except in sediment from deeper layers (>5cm) of the M station. While there are two ways to interpret the lack of methane build up in the batch incubations, i.e., either the absence of methane production or the presence of simultaneous methane production and consumption, our radiotracer data point towards the latter scenario. Together, the radiotracer and batch incubations strongly suggest simultaneous methane production and consumption by the cryptic methane cycle across the salinity transect.

### 4.2. Spatial geochemical trends and electron acceptors for AOM

Another goal of this study was to compare porewater salinity and electron acceptor availability with metabolic processes (i.e., sulfate reduction and AOM) along the transect. Although the salinity in the overlying water indicated a natural salinity gradient within the CSMR due to freshwater and seawater convergence, the porewater salinity (max 73 and 47 PSU at BL and BH, respectively) and sulfate concentrations (max 62 and 39 mM at BL and BH, respectively) in the brackish stations increased with increasing sediment depth and were considerably higher than expected (Fig 2D and J.). This finding suggests that the sediment hydrology of the CSMR is heavily influenced by seawater infiltration rather than by freshwater sources (i.e., by watersheds located more inland). Alternatively, it may also suggest that the CSMR was once a hypersaline coastal wetland. The abundance of salt within salt marshes such as the CSMR is largely controlled by tidal infiltration and inundation and by evaporation (Gardner, 2007; Li et al., 2023; Moffett et al., 2010). For the CSMR sediments to have accumulated such high amounts of salts, the area must have had periodic inundation by seawater to deliver the salt, followed by evaporation. It is conceivable that subsurface hydrology and the geomorphology of the CSMR plays a role in the salinization and/or dilution of the salt in the subsurface of the CSMR (Reddy et al., 2022). However, the present study did not determine the direction of seawater penetration/infiltration (vertical or lateral diffusion) but future studies at the CSMR and other coastal wetlands would benefit from such determinations because the seawater delivers sulfate, which is a key electron acceptor for AOM.

Across the CSMR salinity gradient, sediment porewater sulfate was always high (≥9 mM, Fig. 2 D, J, P, V), with the exception of the 0-1 cm section at the BH station (0.15 mM). Accordingly, these non-limiting sulfate concentrations support high sulfate reduction activity in the top 5-10 cm of the sediment. Sulfate further strongly correlated with microbial communities at the HP station (Fig. 4A and B), which had the highest sulfate reduction rates (Fig. 2V). Notably, while a part of the sulfate reduction was likely linked to organic matter degradation, another part could be coupled to AOM in layers where both processes coincided (e.g., 1-6, 1-10, 1-18, and 13-15 cm sections at the BL, BH, M, and HP stations, respectively) (Fig. 2D, E, J, K, P, Q, V, and W). In support of this hypothesis, 16S sequencing data confirmed the presence of bacterial groups capable of sulfate reduction (Fig. 7C and D) in all layers where ANME groups were found (Fig. 6C and D) and where sulfate reduction and AOM activity overlapped (Fig. 2D, J, P, and V), except at the BL and BH stations, where ANME groups were not detected (see section 4.3 for details). High concentrations of porewater sulfate along the CSMR salinity transect, despite high sulfate reduction activity, suggests that sediment within salt marshes like the CSMR are sources of electron acceptors, which are important to buffer methane production by competitive methanogenesis pathways and support methane consumption by AOM.

Porewater iron (II) and sulfide concentrations (Fig. 2 C, I, O, and U) revealed that sediment below the top 2 cm in the CSMR transitioned from a predominantly iron-reducing (high Fe (II), absence of sulfide) environment to a predominantly sulfate-reducing (low Fe (II), presence of sulfide) environment between the BL and the HP stations. In accordance, the microbial community analysis shows a strong correlation between high iron (II) concentrations and microbial communities at the BL and BH stations suggesting the presence of groups that are involved in iron cycling (Fig. 4B). The 16S rRNA sequencing recovered members of the Acidobacteriota, Firmicutes, Deferrisomatota, and Sva 0485 phyla, which are known to contain lineages of iron-reducing bacteria (Fig. 7A, B, C, and D) (Weber et al., 2006). Additionally, we also found that in the iron-reducing layers in the BL and BH stations were dominated by members of the Desulfatiglandales, Desulfobacterales, and Desulfobulbales, bacterial orders that also have iron-reducing members and may be contributing to the buildup of iron (II) at these stations (Fig. 7A and B).

At the M and HP stations, porewater sulfide was more abundant than at the BL and BH stations (Fig. 2O and U). The sulfide production is likely attributed to the widespread sulfate reduction activity at both stations, and sulfate reduction coupled to AOM (Fig. 2P and V). These findings fit with the microbial community data, which show that microbial communities at the M station are correlated with sulfide and AOM activity, while at the HP station communities are correlated with sulfide and sulfate reduction rates (Fig. 4A and B). At the M station, high concentrations of porewater iron (II) (Fig. 2O) further coincided with microbial communities capable of iron reduction (Fig. 4B and 7C). The 16S rRNA gene diversity at the M station shows a mixture of putative iron-reducing bacteria (i.e., Acidobacteriota, Firmicutes, Geobacterales, Geopsychrobacteraceae, Geothermobacteraceae, and Desulfuromondaceaae) and sulfate/iron-reducing bacteria (i.e., Desulfobacterales and Desulfobulbales) throughout the sediment (Fig. 6C and 7C). This mixture of microbial groups could potentially be contributing to the large buildup of both iron (II) and sulfide in the sediment at the M station (Fig. 2O). Whereas, at the HP station, sulfate/iron-reducing bacteria (i.e., Desulfobacterales and Desulfobulbales) that were detected near the sediment surface (Fig. 7D) could potentially be responsible for the high sulfate reduction activity and high sulfide concentrations near the sediment surface (Fig. 2U and V). The 16S rRNA sequencing data also detected an iron reducer (i.e., Geothermobacteraceae) and sulfate/iron reducers (i.e., Desulfobacterales and Desulfobulbales) (Fig. 7D) just below the sulfate-reducing layer which potentially is responsible for the small amount of iron (II) buildup (Fig. 2U).

The geochemical and molecular data in this study strongly indicate that both iron reduction and sulfate reduction are concurrently active, potentially by groups of iron-reducing and sulfate-reducing bacteria that are present at the M station (Fig 2O and 7C), and to a lesser extent at the HP station (Fig. 2U and 7D). Simultaneous iron and sulfate reduction has been observed in marine (Postma and Jakobsen, 1996), lacustrine (Motelica-Heino et al., 2003) and wetland sediment (Reddy and DeLaune, 2008). The detection of simultaneous iron and sulfate reduction activity is interesting since iron reduction is thermodynamically more favorable than sulfate reduction for shared substrates such as hydrogen and acetate (Jørgensen, 2000) and would typically suppress sulfate reduction activity and ban it to deeper sediment intervals (Lovley and Phillips, 1987). However, iron oxides have been shown to persist below sulfate reduction layers in sedimentary environments that receive high inputs of iron-oxides and/or have sediments that have undergone transient diagenesis (Egger et al., 2015).

In summary, our study shows that CSMR sediments are rich in sulfate and iron (Fig. 2C, D, I, J, O, P, U, V), supporting organoclastic sulfate- and iron-reducing communities (Fig. 7). These sediments thereby not only regulate methane emissions by suppressing competitive methanogenesis but also support AOM, as part of the cryptic methane cycle, linked to potentially both sulfate and iron reduction (Fig. 2E, K, Q, W).

Iron-dependent AOM and has been shown to occur in marine (Beal et al., 2009; Rooze et al., 2016; Schnakenberg et al., 2021), brackish (Egger et al., 2017; Egger et al., 2015), and freshwater (Bar-Or et al., 2017; Ettwig et al., 2016; Leu et al., 2020; Martinez-Cruz et al., 2018), and coastal wetland environments (Segarra et al., 2013; Valenzuela et al., 2019; Wallenius et al., 2021). Iron (III) could be an important electron acceptor for AOM in deeper sediment intervals at the BL, BH, and M stations, where iron (II) is high (Fig. 2C, P and I). However, ANME lineages that are known to conduct iron-dependent AOM were not detected at the BL and BH stations (Fig. 6A and B). At the M station, ASVs belonging to ANME 2-a and -c (Fig. 6C), which have recently been demonstrated to be capable of iron-dependent AOM in marine sediments (Aromokeye et al., 2020; Scheller et al., 2016), were found. Although the present study did not directly test for the coupling of iron to AOM, our study shows a substantial buildup of iron (II) in the sediment at BL, BH and M stations (Fig. 2C, I, and O) and a slight buildup of iron (II) in the presence of sulfide a the HP station (Fig. 2U) indicating that iron (III) could potentially be an important electron acceptor for AOM. Recent work at the CSMR does strongly suggest that both sulfate and iron (III) are important electron acceptors for AOM in the HP station at the CSMR (Liu, 2024). However, more work is needed to better connect the role of iron and associated microbial communities to the cryptic methane cycle within the CSMR.

### 4.3. Archaeal methanogen and methanotroph communities

Another goal of this study was to elucidate the potential archaea lineages responsible for methanogenesis and AOM activity across the CSMR transect and to investigate whether they cooccur in the same sediment horizons, consistent with the potential for cryptic methane cycling in the CSMR. At the BL and BH stations, MG-MMA activity was strongly correlated with microbial community composition (Fig. 4A). At the BL and BH stations, low abundances of *Methanosarcinaceae* (Fig. 6A and B), a methanogenic family capable of producing methane from non-competitive substrates (Boone et al., 2015), overlapped with methylotrophic methanogenesis (Fig. 2E and K). Bathyarchaeia, detected at the BL and BH stations, may also contribute to the methylotrophic methanogenesis activity observed there. Metabolism reconstructions from metagenomes revealed that some groups of Bathyarchaeia have genes encoding for the methyl-coenzyme M reductase complex and are potentially capable of both methylotrophic methanogenesis and methane-oxidation (Evans et al., 2015; Hou et al., 2023). However, this archaeal group is highly diverse and the evidence presented in this study does not directly link the methanogenesis activity to Bathyarchaeia. Future studies should investigate the potential contribution of these groups to the global methane budget.

At the M station, the relative abundance of ANME 2a-c ASVs dominated over methanogen lineages in shallower sediment intervals (< 9 cm) (Fig. 5C) and were likely responsible for the detected AOM-CH_4_ activity, maintaining low environmental methane concentrations in these horizons (Fig 2 and R). ASVs belonging to ANME 2a-2c were also found in the 12.5-13.5 cm horizon, which was consistent with the dramatic increase in AOM activity in this horizon and the decline in methane above and below it (Fig. 2Q and R). In this horizon, ASVs assigned to methylotrophic and acetoclastic methanogens (families *Methanosarcinaceae, Methanoregulaceae*, and *Methanosaetaceae)* were detected, suggesting that methanogens in the M station are capable of acetoclastic and non-competitive methanogenesis and ANME are coexisting in sediment horizons where sulfate reduction was active (Fig. 2P). ASVs belonging to ANMEs were not detected between 13.5 and 19.5 cm despite the measurement of higher AOM activity (Fig. 2Q). Instead, ASVs belonging to *Methanosarcinaceae*, *Methanoregulaceae*, and *Methanosaetaceae* extended into these deeper horizons, which could be linked to the higher methane concentrations found in these depths (Fig. 2R) as well as the higher levels of methane temporarily produced in methanogenesis batch incubations with deeper sections (Fig. 3C).

At the HP station, ANME 2a-2c families were found in the top 2.5 cm even though no AOM activity was detected by ^14^C-CH_4_ incubations in this layer. As the 16S data does not provide direct evidence of metabolic activity, it is possible that the ANMEs were present but not active, potentially due to the high salinity in the HP station (Fig. 2V), causing osmotic stress and lowering or pausing AOM activity (Oren, 2011). In deeper sediment intervals, two methanogenic families belonging to *Methanosarcinaceae* and *Methanonatronarchaeaceae* were detected. Both of these groups are halotolerant and can produce methane from non-competitive substrates and could be responsible for the MG-MMA rates we detected in the deeper sediments at the HP station (Boone et al., 2015; Sorokin et al., 2018).

An alternative hypothesis worth testing in future investigations is that methanogens are potentially responsible for mediating both methanogenesis and AOM in the cryptic methane cycle. Methanogens and ANMEs both carry the Mcr gene that encodes for the methyl coenzyme M, which depending on the thermodynamic conditions can work preferentially in either the reductive or oxidative directions (Hallam et al., 2003; Holler et al., 2011; Timmers et al., 2017). In addition, methanogenic groups within the *Methanosarcinales* order, which were detected in the CSMR sediments (Fig. 6) have been implicated in performing AOM coupled to iron reduction of poorly reactive iron oxides in lake sediment (Bar-Or et al., 2017) and in marine sediment (Liang et al., 2019; Yan et al., 2023; Yan et al., 2018). Testing the involvement of members of the *Methanosarcinales* in iron-dependent AOM could be particularly interesting for sediment intervals in which we detected simultaneous AOM activity (Fig. 2E, K, and Q), high porewater iron (II) concentrations (Fig. C, I, and O) and ASVs belonging to *Methanosarcinales* (Fig. 6A, B, C, and D). At the HP station, AOM was detected in deeper sediment intervals where no ANME ASVs were recovered. Here, AOM may also be mediated by groups of methanogens. At 16.5-17.5 cm, *Methanosarcinaceae*, a group within *Methanosarcinales*, was detected and could potentially contributing to the measured rates of AOM similar to the other stations. However, iron (II) concentrations at the HP station were low and it is unclear if iron (III) could be the electron acceptor for AOM at these depths. In combination, our data point to a potential involvement of methanogens in iron-dependent methane oxidation. Future investigations should focus on deciphering the mechanisms methanogens might be using to oxidize methane with iron as the electron acceptor and whether this potential catabolic reaction is coupled to growth of these archaea.

### 4.4. Indications for rapid turnover of metabolic substrates

Salt marshes are known to be rich in fresh organic matter whose breakdown intermediates provide competitive and non-competitive substrates for methanogenesis (Fitzsimons et al., 2005; Fitzsimons et al., 1997; Wang and Lee, 1994; Yuan et al., 2014; 2019). Porewater analysis by NMR revealed that porewaters from the BL, BH, and M stations had quantifiable concentrations of acetate. Acetate is a product from fermentation of organic matter by homoacetogenesis (Jørgensen, 2000; Ragsdale and Pierce, 2008). Acetate is an important substrate for a wide range of microbial groups and considered a competitive substrate between methanogenic archaea and sulfate-reducing bacteria (Conrad, 2020; Jørgensen, 2000). Acetate concentrations that were found to be below quantification or detection at our studied stations could point to rapid metabolic turnover, similar to what has been described for hydrogen (Conrad, 1999; Hoehler et al., 2001).

Porewater methanol was mostly below detection and sometimes present but not quantifiable in the CSMR sediment porewater (see section 3.1.4.). Although methanol is known to be a non-competitive substrate for methylotrophic methanogenesis in coastal wetlands (King et al., 1983; Oremland and Polcin, 1982) it has recently been found to be a carbon source for other non-methanogenic anaerobic methylotrophs such as denitrifying and sulfate-reducing (Fischer et al., 2021), which according to our 16S data are likely present in the CSMR (Fig. 6 and 7). As this substrate is expected to be produced in surface sediments of salt marshes (Oremland et al., 1982; Oremland and Polcin, 1982), low or undetectable concentrations of methanol in the porewater could indicate fast metabolic turnover, similar to acetate.

Natural porewater MMA concentrations were also mostly below detection (<3 µM), however, at some depth intervals in the BL and HP stations MMA concentrations were below quantification (<10 µM), but detectable (Table 1). It is not possible to report definitive quantities of MMA in this study, however, we can bracket the MMA concentrations in a range between 3 and 10 µM. MMA concentrations in sediment porewaters are still poorly constrained. A previous study of MMA in the Mersey Estuary, UK reports sediment porewater concentrations up to 319 µM (Fitzsimons et al., 1997). Other studies report lower sediment porewater concentrations, for example ∼2 µM MMA in the Flax Pond salt marsh (Wang and Lee, 1994) and between 0.08 and 1.44 µM in the Thames Estuary, UK (Fitzsimons et al., 2006). Low MMA concentrations in the CSMR sediment porewater may as well indicate rapid metabolic consumption by the microbial community and/or binding to mineral surfaces (Wang and Lee, 1990; Xiao et al., 2022). Data from our ^14^C-MMA incubations provide support for both hypotheses as we observed the metabolic potential for MMA through the turnover of the injected ^14^C-MMA and found that between 38% and 55% of the injected ^14^C-MMA bound to the sediment (see Sec. 3.3.1).

Interestingly, we found large differences between AOM-CH_4_ (i.e., AOM determined based on ^14^C-TIC production after injection of ^14^C-CH_4_) and AOM-MMA (AOM determined based on ^14^C-TIC production after injection of ^14^C-MMA) at all CSMR stations. In intervals where both AOM-CH_4_ and AOM-MMA rates were detected, AOM-MMA was 1-2 orders of magnitude higher than AOM-CH_4_ (Fig. 2E, K, Q, and W). We suggest the large difference between the two AOM rates is the result of ^14^C-TIC production from direct oxidation of ^14^C-MMA by non-methanogenic pathways, similarly to what was previously hypothesized by (Krause et al., 2023). Direct conversion of ^14^C-MMA to ^14^C-TIC would incorrectly inflate the rate constants for AOM-MMA dramatically (see Eq. 2). Figure 2F, L, R, and X indeed show that rate constants for AOM-MMA are 1-2 orders of magnitude higher than both MG-MMA and AOM-CH_4_. We therefore suggest that the ^14^C-TIC produced during the ^14^C-MMA incubations stems only partially from AOM as part of the cryptic methane cycle (i.e., via the ^14^C-CH_4_ intermediate). The majority of ^14^C-MMA was likely subject to direct methylamine oxidation by an unidentified anaerobic methylotrophic metabolism.

Methylamines are the simplest alkylated amine and derived from the degradation of osmolytes found in plant biomass (Oren, 1990; Taubert et al., 2017). Methylamines are ubiquitously found in saline and hypersaline conditions in marine sediments (Mausz and Chen, 2019; Zhuang et al., 2016; Zhuang et al., 2017) and in coastal wetlands (Fitzsimons et al., 2005; Fitzsimons et al., 1997; Fitzsimons et al., 2001; Fitzsimons et al., 2006). Because methylamine molecules are both a carbon and nitrogen source it is an important food sources for a variety of microbial communities such as aerobic methylotrophic bacteria (Chistoserdova, 2015; Hanson and Hanson, 1996; Taubert et al., 2017) and obligate anaerobic methanogens (Chistoserdova, 2015; Thauer, 1998). However, increasing reports are providing evidence that methylamines may be directly oxidized anaerobically by non-methanogenic metabolisms in anoxic sediment (Cadena et al., 2018; De Anda et al., 2021; Farag et al., 2021; Kivenson et al., 2021; Zhuang et al., 2019). Although the present study does not provide direct evidence of non-methanogenic anaerobic methylamine oxidation, based on the geochemical and molecular results in the present study we propose sulfate-reducing bacteria are a potential consumer of MMA in the CSMR in addition to methylotrophic methanogenesis. Kivenson et al., (2021) recently reanalyzed the metagenomes and metatranscriptomes of Desulfobacterales collected from sulfidic sediment in the Baltic Sea (Thureborn et al., 2016) and the Columbia River Estuary (Smith et al., 2015; Smith et al., 2019) and found the expression of trimethylamine metabolisms in the Desulfobacterales which strongly suggests that sulfate reducers could be actively in competition with methylotrophic methanogens. As ASVs belonging to the order Desulfobacterales were detected throughout the sediment intervals across the CSMR transect (Fig. 6 and 7), it is possible that members of this group were actively involved in anaerobic methylamine consumption. Future research should investigate with transcriptomic studies to assess if sulfate-reducing lineages in the CSMR sediment are expressing the metabolic machinery required to utilize methylamines.

Our 16S rRNA data also revealed other archaeal and bacterial candidates that could potentially be implicated in methylamine consumption in the CSMR. At the BL and BH station, we detected ASV’s of both Bathyarchaeia and Lokiarchaeia (Fig. 5A and B). Metabolic reconstructions from metagenomes found that members of both groups have the potential for non-methanogenic anaerobic turnover of C1 compounds such as methanol and methylamines (Hou et al., 2023; Sun et al., 2021). Potential bacterial candidates for anaerobic methylamine oxidation in the CSMR transect are Proteobacteria, Firmicutes, Actinobacteria, Verrucomicrobia, candidate phylum NC10, and Actinobacteriota (Fig. 5E-H) as these phyla comprise known methylotrophs (Anthony, 1982; Chistoserdova and Lidstrom, 2013; McTaggart et al., 2015; Zemskaya et al., 2021).

If methylated substrates in anaerobic metabolism are not restricted to putative methanogenic archaea, then it is crucial to understand how competitive these substrates are between methylotrophic methanogens and other microorganisms. A future hypothesis worth testing is that sulfate-reducing methylotrophs limit the availability of methylated substrates to methylotrophic methanogenesis in coastal wetland sediment. If future investigations support this hypothesis, this process will contribute to the comparably low methane emissions from coastal wetlands to the atmosphere and thus better constrain the global methane budget. Identifying this unknown metabolism, the responsible microbial groups, and the competition mechanism is crucial. If environmental conditions shift to favor methanogenic archaea, it remains unclear whether the existing cryptic methane cycle will be amplified or if coastal wetlands will become a larger methane source.

### 4.5. Implications for cryptic methane cycling in coastal wetlands

Coastal wetlands are at the boundary between terrestrial and marine ecosystems. However, reports showing the decline in coastal wetland areas due to a variety of anthropogenic pressures (i.e., land reclamation, agriculture, and run off) are increasing (Newton et al., 2020). The leading concern is sea-level rise caused by increasing temperatures globally driven by climate change. The fear is that sea level rise will potentially permanently inundate coastal wetland areas globally. This inundation of seawater would cause changes to biogeochemical cycles and deliver more organic material into coastal wetlands, leading to changes in biogeochemical cycles and promoting methanogenesis in coastal wetland sediment (Chambers et al., 2013; Dinsmore et al., 2009; Gatland et al., 2014; Liu et al., 2019; Vizza et al., 2017; Zhao et al., 2020). With this imminent problem arising, important questions remain about the effect of climate-induced sea-level rise has on methylotrophic activity and the cryptic methane cycle within coastal wetlands. A future hypothesis worth testing is that inundation of coastal wetlands due to sea-level rise enhances cryptic methane cycling activity (elevated methanogenesis and AOM activity) supported by large pools of electron acceptors found in the subsurface porewaters. If confirmed, with the evidence presented in this study, the cryptic methane cycle maybe enhanced during periods of extended inundation, keeping methane concentrations low and therefore a lower emission of a potent greenhouse gas.

## 5. Conclusions

In the present study, we set about to find spatial evidence of cryptic methane cycling and the potential microbial communities involved, in the surface sediments along a land-ocean, 4-station transect within a southern Californian salt marsh. We found spatial overlap of methylotrophic methanogenesis and AOM activity in different depth horizons of all four stations. We conclude that the cryptic methane cycle is active keeping methane concentrations present but low across the salt marsh transect. We further found that the salinity at the sediment-water interface of the supposedly brackish stations was different from the subsurface, where high salinity and high availability of sulfate prevailed, suggesting that more inland portions of the saltmarsh may have once been more hypersaline. At none of the four stations sulfate was limiting in the subsurface, supporting sulfate reduction activity and likely linked AOM. However, the concomitant presence of porewater iron (II) and ASV’s belonging to heterotrophic iron-reducing bacteria and ANME known to couple methane oxidation with iron reduction indicate that iron reduction could, too, be an important metabolism and potentially be coupled to AOM. Metabolomic analysis of porewater indicate a rapid production and consumption of methanogenic substrates, including acetate, methanol, and MMA. While molecular data revealed methanogenic archaea at all stations, ANME were present only at the marine and hypersaline stations. This finding suggest that cryptic methane cycling could be facilitated by either a methanogen-methanotroph archaea couple, where both groups coincided, or by putative methanogenic archaea capable of both processes, where ANME were absent. Our data from radioisotope incubations strongly suggest that not only is cryptic methane cycling active in the salt marsh sediment, but there is also an unknown anaerobic methylotrophic metabolism directly oxidizing methylamine into the inorganic carbon pool. Our molecular and biogeochemical analyses along with literature evidence identified sulfate-reducing bacteria as a potential candidate responsible for the turnover of methylated substrates, but more work is needed to confirm. Our study emphasizes the biogeochemical complexity occurring in the sediment across spatial gradients within a salt marsh. Future work should investigate the mechanisms and dynamics of the production and consumption of methylated substrates like methylamine and how they may affect the cryptic methane cycle. If methylated substrates are indeed another competitive substrate between methanogens and sulfate reducers, such investigations may uncover additional reasons for low methane emissions from salt marshes.

## Acknowledgements

The authors thank the University of California Natural Reserve System and the Project Scientist of the Carpinteria Salt Marsh Reserve, A. Brooks, for authorizing the field sampling in June 2019. We acknowledge X. Hwang for providing fieldwork and laboratory instrument support. We acknowledge S. Connon and S. Lim for providing laboratory and sequencing analysis support. This work received financial support through the University of California and the National Science Foundation (NSF) (Award No.: 1852912). The NMR experiments in this study were performed under a limited scope project awarded (proposal 51757: 10.46936/ltds.proj.2020.51757/60006896) to T. Treude using the Environmental Molecular Sciences Laboratory, a DOE Office of Science User Facility sponsored by the Biological and Environmental Research program under Contract No. DE-AC05-76RL01830. V. J. Orphan and R. Wipfler contributions were supported through the U.S. Department of Energy, Office of Science, Office of Biological and Environmental Research under Award Number DE-SC0022991.

## Data Availability

Datasets from geochemical and ^14^C analysis are accessible via the Biological & Chemical Oceanography Data Management Office (BCO-DMO) through the following DOI [# will be provided before publication]. Data from the 16S rRNA molecular analysis are accessible via the National Center for Biotechnology Information (NCBI) BioProject PRJNA1199032 through the following DOI [https://www.ncbi.nlm.nih.gov/bioproject/PRJNA1199032].

## CRediT authorship contribution statement

**Sebastian J. E. Krause:** Writing-Original Draft, Conceptualizing, Methodology, Validation, Formal analysis, Investigation, Data curation, Visualization, Supervision, and Project administration. **Rebecca Wipfler:** Methodology, Visualization, Software, Validation, Formal analysis, Data curation, Writing-review and editing. **Jiarui Liu:** Investigation, Writing- review and editing. **David J. Yousavich:** Investigation, Writing- review and editing. **DeMarcus Robinson:** Investigation, Writing- review and editing. **David W. Hoyt:** Investigation, Resources, Formal analysis, Writing- review and editing. **Victoria J. Orphan:** Investigation, Resources, Formal analysis, Supervision, Writing- review and editing. **Tina Treude:** Conceptualization, Methodology, Validation, Resources, Data curation, Writing-review and editing, Supervision, Project Administration, Funding acquisition.

## References

Anthony, C., 1982. The biochemistry of methylotrophs.

Aromokeye, D.A., Kulkarni, A.C., Elvert, M., Wegener, G., Henkel, S., Coffinet, S., Eickhorst, T., Oni, O.E., Richter-Heitmann, T., Schnakenberg, A., 2020. Rates and microbial players of iron-driven anaerobic oxidation of methane in methanic marine sediments. Frontiers in Microbiology 10, 487993.

Bar-Or, I., Elvert, M., Eckert, W., Kushmaro, A., Vigderovich, H., Zhu, Q., Ben-Dov, E., Sivan, O., 2017. Iron-coupled anaerobic oxidation of methane performed by a mixed bacterial-archaeal community based on poorly reactive minerals. Environmental science & technology 51(21), 12293–12301.

Barnes, R.O., Goldberg, E.D., 1976. Methane production and consumption in anoxic marine sediments. Geology 4, 297–300.

Bartlett, K.B., Bartlett, D.S., Harriss, R.C., Sebacher, D.J., 1987. Methane emissions along a salt marsh salinity gradient. Biogeochemistry 4, 183–202.

Bartoli, M., Nizzoli, D., Zilius, M., Bresciani, M., Pusceddu, A., Bianchelli, S., Sundbäck, K., Razinkovas-Baziukas, A., Viaroli, P., 2021. Denitrification, nitrogen uptake, and organic matter quality undergo different seasonality in sandy and muddy sediments of a turbid estuary. Frontiers in microbiology 11, 612700.

Beal, E.J., House, C.H., Orphan, V.J., 2009. Manganese- and iron-dependent marine methane oxidation. Science 325, 184–187.

Bernard, B.B., 1979. Methane in marine sediments. Deep Sea Research Part A. Oceanographic Research Papers 26(4), 429–443.

Beulig, F., Røy, H., McGlynn, S.E., Jørgensen, B.B., 2018. Cryptic CH4 cycling in the sulfate-methane transition of marine sediments apparently mediated by ANME-1 archaea. The ISME journal, 10.1038/s41396-41018-40273-z.

Boetius, A., Ravenschlag, K., Schubert, C.J., Rickert, D., Widdel, F., Giesecke, A., Amann, R., Jørgensen, B.B., Witte, U., Pfannkuche, O., 2000. A marine microbial consortium apparently mediating anaerobic oxidation of methane. Nature 407, 623–626.

Booij, K., Helder, W., Sundby, B., 1991. Rapid redistribution of oxygen in a sandy sediment induced by changes in the flow velocity of the overlying water. Netherlands Journal of Sea Research 28(3), 149–165.

Boone, D.R., Whitman, W.B., Koga, Y., 2015. Methanosarcinaceae. Bergey’s Manual of Systematics of Archaea and Bacteria, 1–2.

Boudreau, B.P., Huettel, M., Forster, S., Jahnke, R.A., McLachlan, A., Middelburg, J.J., Nielsen, P., Sansone, F., Taghon, G., Van Raaphorst, W., 2001. Permeable marine sediments: overturning an old paradigm. EOS, Transactions American Geophysical Union 82(11), 133–136.

Bridgham, S., Cadillo-Quiroz, H., Keller, J., Zhuang, Q., 2013. Methane emissions from wetlands: biogeochemical, microbial, and modeling perspectives from local to global scales. Global Change Biology 19(5), 1325–1346. doi.org/10.1111/gcb.12131.

Bridgham, S., Megonigal, J., Keller, J., Bliss, N., Trettin, C., 2006. The carbon balance of North American wetlands. Wetlands 26(4), 889–916. doi.org/10.1672/0277-5212(2006)26[889:TCBONA]2.0.CO;2.

Cadena, S., García-Maldonado, J.Q., López-Lozano, N.E., Cervantes, F.J., 2018. Methanogenic and sulfate-reducing activities in a hypersaline microbial mat and associated microbial diversity. Microbial ecology 75(4), 930–940.

Callahan, B.J., McMurdie, P., Rosen, M., Han, A., Johnson, A., 2016. JA, Holmes SP. 2016. DADA2: high-resolution sample inference from Illumina amplicon data. Nature Methods 13(7), 581–583.

Cao, M., Xin, P., Jin, G., Li, L., 2012. A field study on groundwater dynamics in a salt marsh–Chongming Dongtan wetland. Ecological Engineering 40, 61–69.

Carol, E.S., Dragani, W.C., Kruse, E.E., Pousa, J.L., 2012. Surface water and groundwater characteristics in the wetlands of the Ajó River (Argentina). Continental Shelf Research 49, 25–33.

Carol, E.S., Kruse, E.E., Pousa, J.L., 2011. Influence of the geologic and geomorphologic characteristics and of crab burrows on the interrelation between surface water and groundwater in an estuarine coastal wetland. Journal of Hydrology 403(3-4), 234–241.

Chadwick, G.L., Skennerton, C.T., Laso-Pérez, R., Leu, A.O., Speth, D.R., Yu, H., Morgan-Lang, C., Hatzenpichler, R., Goudeau, D., Malmstrom, R., 2022. Comparative genomics reveals electron transfer and syntrophic mechanisms differentiating methanotrophic and methanogenic archaea. PLoS biology 20(1), e3001508.

Chambers, L.G., Osborne, T.Z., Reddy, K.R., 2013. Effect of salinity-altering pulsing events on soil organic carbon loss along an intertidal wetland gradient: a laboratory experiment. Biogeochemistry 115, 363–383.

Chen, S., Torres, R., Goñi, M.A., 2016. The role of salt marsh structure in the distribution of surface sedimentary organic matter. Estuaries and coasts 39, 108–122.

Chistoserdova, L., 2015. Methylotrophs in natural habitats: current insights through metagenomics. Applied microbiology and biotechnology 99(14), 5763–5779.

Chistoserdova, L., Lidstrom, M.E., 2013. Aerobic methylotrophic prokaryotes. The prokaryotes, 267–285.

Cline, J.D., 1969. Spectrophometric determination of hydrogen sulfide in natural waters. Limnol. Oceanogr. 14, 454–458.

Conrad, R., 1999. Contribution of hydrogen to methane production and control of hydrogen concentrations in methanogenic soils and sediments. FEMS microbiology Ecology 28(3), 193–202.

Conrad, R., 2020. Importance of hydrogenotrophic, aceticlastic and methylotrophic methanogenesis for methane production in terrestrial, aquatic and other anoxic environments: a mini review. Pedosphere 30(1), 25–39.

Dale, A.W., Sommer, S., Lomnitz, U., Montes, I., Treude, T., Liebetrau, V., Gier, J., Hensen, C., Dengler, M., Stolpovsky, K., Bryant, L.D., Wallmann, K., 2015. Organic carbon production, mineralisation and preservation on the Peruvian margin. Biogeosciences 12, 1537–1559.

De Anda, V., Chen, L.-X., Dombrowski, N., Hua, Z.-S., Jiang, H.-C., Banfield, J.F., Li, W.-J., Baker, B.J., 2021. Brockarchaeota, a novel archaeal phylum with unique and versatile carbon cycling pathways. Nature communications 12(1), 1–12.

de los Santos, C.B., Egea, L.G., Martins, M., Santos, R., Masqué, P., Peralta, G., Brun, F.G., Jiménez-Ramos, R., 2023. Sedimentary organic carbon and nitrogen sequestration across a vertical gradient on a temperate wetland seascape including salt marshes, seagrass meadows and rhizophytic macroalgae beds. Ecosystems 26(4), 826–842.

del Pilar Alvarez, M., Carol, E., Hernández, M.A., Bouza, P.J., 2015. Groundwater dynamic, temperature and salinity response to the tide in Patagonian marshes: Observations on a coastal wetland in San José Gulf, Argentina. Journal of South American Earth Sciences 62, 1–11.

Diao, M., Dyksma, S., Koeksoy, E., Ngugi, D.K., Anantharaman, K., Loy, A., Pester, M., 2023. Global diversity and inferred ecophysiology of microorganisms with the potential for dissimilatory sulfate/sulfite reduction. FEMS Microbiology Reviews 47(5), fuad058.

Dinsmore, K.J., Skiba, U.M., Billett, M.F., Rees, R.M., 2009. Effect of water table on greenhouse gas emissions from peatland mesocosms. Plant and Soil 318, 229–242.

Egger, M., Hagens, M., Sapart, C.J., Dijkstra, N., van Helmond, N.A., Mogollón, J.M., Risgaard-Petersen, N., van der Veen, C., Kasten, S., Riedinger, N., 2017. Iron oxide reduction in methane-rich deep Baltic Sea sediments. Geochimica et Cosmochimica Acta 207, 256–276.

Egger, M., Rasigraf, O., Sapart, C.J., Jilbert, T., Jetten, M.S., Rockmann, T., Van der Veen, C., Banda, N., Kartal, B., Ettwig, K.F., 2015. Iron-mediated anaerobic oxidation of methane in brackish coastal sediments. Environmental Science & Technology 49(1), 277–283.

Ettwig, K.F., Zhu, B., Speth, D., Keltjens, J.T., Jetten, M.S., Kartal, B., 2016. Archaea catalyze iron-dependent anaerobic oxidation of methane. Proceedings of the National Academy of Sciences 113(45), 12792–12796.

Evans, P.N., Parks, D.H., Chadwick, G.L., Robbins, S.J., Orphan, V.J., Golding, S.D., Tyson, G.W., 2015. Methane metabolism in the archaeal phylum Bathyarchaeota revealed by genome-centric metagenomics. Science 350(6259), 434–438.

Farag, I.F., Zhao, R., Biddle, J.F., 2021. “Sifarchaeota,” a Novel Asgard Phylum from Costa Rican Sediment Capable of Polysaccharide Degradation and Anaerobic Methylotrophy. Applied and environmental microbiology 87(9), e02584–02520.

Fischer, P.Q., Sánchez-Andrea, I., Stams, A.J., Villanueva, L., Sousa, D.Z., 2021. Anaerobic microbial methanol conversion in marine sediments. Environmental microbiology 23(3), 1348–1362.

Fitzsimons, M., Dawit, M., Revitt, D., Rocha, C., 2005. Effects of early tidal inundation on the cycling of methylamines in inter-tidal sediments. Marine Ecology Progress Series 294, 51–61.

Fitzsimons, M.F., Jemmett, A.W., Wolff, G.A., 1997. A preliminary study of the geochemistry of methylamines in a salt marsh. Organic geochemistry 27(1-2), 15–24.

Fitzsimons, M.F., Kahni-danon, B., Dawitt, M., 2001. Distributions and adsorption of the methylamines in the inter-tidal sediments of an East Anglian Estuary. Environm. Experiment. Bot. 46, 225–236.

Fitzsimons, M.F., Millward, G.E., Revitt, D.M., Dawit, M.D., 2006. Desorption kinetics of ammonium and methylamines from estuarine sediments: consequences for the cycling of nitrogen. Marine Chemistry 101(1-2), 12–26.

Gardner, L.R., 2007. Role of stratigraphy in governing pore water seepage from salt marsh sediments. Water Resources Research 43(7).

Gatland, J.R., Santos, I.R., Maher, D.T., Duncan, T., Erler, D.V., 2014. Carbon dioxide and methane emissions from an artificially drained coastal wetland during a flood: Implications for wetland global warming potential. Journal of Geophysical Research: Biogeosciences 119(8), 1698–1716.

Goslin, J., Sansjofre, P., Van Vliet-Lanoë, B., Delacourt, C., 2017. Carbon stable isotope (δ13C) and elemental (TOC, TN) geochemistry in saltmarsh surface sediments (Western Brittany, France): a useful tool for reconstructing Holocene relative sea-level. Journal of Quaternary Science 32(7), 989–1007.

Grasshoff, K., Ehrhardt, M., Kremling, K., 1999. Methods of seawater analysis. Wiley-VCH Verlag GmbH, Weinheim.

Hallam, S.J., Girguis, P.R., Preston, C.M., Richardson, P.M., DeLong, E.F., 2003. Identification of methyl coenzym m reductase A (mcrA) genes associated with methane-oxidizing archaea. Appl. Environ. Microbiol. 69(9), 5483–5491.

Hanson, R.S., Hanson, T.E., 1996. Methanotrophic bacteria. Microbiol. Rev. 60(2), 439–471.

Hinrichs, K.-U., Boetius, A., 2002. The anaerobic oxidation of methane: new insights in microbial ecology and biogeochemistry, in: Wefer, G., Billett, D., Hebbeln, D., Jørgensen, B.B., Schlüter, M., Van Weering, T. (Eds.), Ocean Margin Systems. Springer-Verlag, Berlin, pp. 457–477.

Hoehler, T.M., Alperin, M.J., Albert, D.B., Martens, C.S., 2001. Apparent minimum free energy requirements for methanogenic Archaea and sulfate-reducing bacteria in an anoxic marine sediment. FEMS Microbiol. Ecol. 38, 33–41.

Holler, T., Wegener, G., Niemann, H., Deusner, C., Ferdelman, T.G., Boetius, A., Brunner, B., Widdel, F., 2011. Carbon and sulfur back flux during anaerobic microbial oxidation of methane and coupled sulfate reduction. Proceedings of the National Academy of Sciences 108(52), E1484–E1490.

Holmes, D.E., Nicoll, J.S., Bond, D.R., Lovley, D.R., 2004. Potential role of a novel psychrotolerant member of the family Geobacteraceae, Geopsychrobacter electrodiphilus gen. nov., sp. nov., in electricity production by a marine sediment fuel cell. Applied and Environmental Microbiology 70(10), 6023–6030.

Hou, J., Wang, Y., Zhu, P., Yang, N., Liang, L., Yu, T., Niu, M., Konhauser, K., Woodcroft, B.J., Wang, F., 2023. Taxonomic and carbon metabolic diversification of Bathyarchaeia during its coevolution history with early Earth surface environment. Science Advances 9(27), eadf5069.

Huettel, M., Ziebis, W., Forster, S., 1996. Flow-induced uptake of particulate matter in permeable sediments. Limnology and Oceanography 41(2), 309–322.

Huettel, M., Ziebis, W., Forster, S., Luther, I.G.W., 1998. Advective transport affecting metal and nutrient distribution and interfacial fluxes in permeable sediments. Geochim. Cosmochim. Acta 62(4), 613–631.

IPCC, C.C., 2014. Mitigation of climate change. Contribution of working group III to the fifth assessment report of the intergovernmental panel on climate change.

Iversen, N., Jørgensen, B.B., 1993. Diffusion coefficients of sulfate and methane in marine sediments: influence of porosity. Geochim. Cosmochim. Acta 57, 571–578.

Jackson, K.L., Whitcraft, C.R., Dillon, J.G., 2014. Diversity of Desulfobacteriaceae and overall activity of sulfate-reducing microorganisms in and around a salt pan in a southern California coastal wetland. Wetlands 34, 969–977.

Joye, S.B., Boetius, A., Orcutt, B.N., Montoya, J.P., Schulz, H.N., Erickson, M.J., Logo, S.K., 2004. The anaerobic oxidation of methane and sulfate reduction in sediments from Gulf of Mexico cold seeps. Chem. Geol. 205, 219–238.

Jørgensen, B.B., 1978. A comparison of methods for the quantification of bacterial sulphate reduction in coastal marine sediments: I. Measurements with radiotracer techniques. Geomicrobiol. J. 1(1), 11–27.

Jørgensen, B.B., 2000. Bacteria and marine biogeochemistry, in: Schulz, H.D., Zabel, M. (Eds.), Marine biogeochemistry. Springer Verlag, Berlin, pp. 173–201.

Kallmeyer, J., Ferdelman, T.G., Weber, A., Fossing, H., Jørgensen, B.B., 2004. A cold chromium distillation procedure for radiolabeled sulfide applied to sulfate reduction measurements. Limnol. Oceanogr. Methods 2, 171–180.

King, G., Klug, M.J., Lovley, D.R., 1983. Metabolism of acetate, methanol, and methylated amines in intertidal sediments of Lowes Cove, Maine. 45(6), 1848–1853.

Kivenson, V., Paul, B.G., Valentine, D.L., 2021. An ecological basis for dual genetic code expansion in marine deltaproteobacteria. Frontiers in microbiology, 1545.

Knittel, K., Boetius, A., 2009. Anaerobic oxidation of methane: progress with an unknown process. Annu. Rev. Microbiol. 63, 311–334.

Knittel, K., Wegener, G., Boetius, A., 2018. Anaerobic methane oxidizers. Microbial Communities Utilizing Hydrocarbons and Lipids: Members, Metagenomics and Ecophysiology/ed. TJ McGenity. Cham: Springer, 1–21.

Knudsen, M., 1901. Hydrographical Tables According to the Measurings of Carl Forch…[et Al.] and with Assistance of Björn-Andersen…[et Al.]. GEC Gad.

Kostka, J.E., Roychoudhury, A., Van Cappellen, P., 2002. Rates and controls of anaerobic microbial respiration across spatial and temporal gradients in saltmarsh sediments. Biogeochemistry 60, 49–76.

Krause, S.J., Liu, J., Yousavich, D.J., Robinson, D., Hoyt, D.W., Qin, Q., Wenzhöfer, F., Janssen, F., Valentine, D.L., Treude, T., 2023. Evidence of cryptic methane cycling and non-methanogenic methylamine consumption in the sulfate-reducing zone of sediment in the Santa Barbara Basin, California. Biogeosciences 20(20), 4377–4390.

Krause, S.J., Treude, T., 2021. Deciphering cryptic methane cycling: Coupling of methylotrophic methanogenesis and anaerobic oxidation of methane in hypersaline coastal wetland sediment. Geochimica et Cosmochimica Acta 302, 160–174.

Kristjansson, J.K., Schönheit, P., Thauer, R.K., 1982. Different Ks values for hydrogen of methanogenic bacteria and sulfate reducing bacteria: an explanation for the apparent inhibition of methanogenesis by sulfate. Arch. Microbiol. 131, 278–282.

La, W., Han, X., Liu, C.-Q., Ding, H., Liu, M., Sun, F., Li, S., Lang, Y., 2022. Sulfate concentrations affect sulfate reduction pathways and methane consumption in coastal wetlands. Water Research 217, 118441.

Leu, A.O., Cai, C., McIlroy, S.J., Southam, G., Orphan, V.J., Yuan, Z., Hu, S., Tyson, G.W., 2020. Anaerobic methane oxidation coupled to manganese reduction by members of the Methanoperedenaceae. The ISME Journal 14(4), 1030–1041.

Li, Y., Wang, D., Chen, Z., Chen, J., Hu, H., Wang, R., 2021. Methane emissions during the tide cycle of a Yangtze Estuary salt marsh. Atmosphere 12(2), 245.

Li, Z., Hodges, B.R., Shen, X., 2023. Modeling hypersalinity caused by evaporation and surface–subsurface exchange in a coastal marsh. Journal of Hydrology 618, 129268.

Liang, L., Wang, Y., Sivan, O., Wang, F., 2019. Metal-dependent anaerobic methane oxidation in marine sediment: Insights from marine settings and other systems. Science China Life Sciences 62, 1287–1295.

Liu, J., 2024. The Biogeochemistry of Methane Cycling and its Clumped Isotope Effects. University of California, Los Angeles.

Liu, L., Wang, D., Chen, S., Yu, Z., Xu, Y., Li, Y., Ge, Z., Chen, Z., 2019. Methane emissions from estuarine coastal wetlands: Implications for global change effect. Soil Science Society of America Journal 83(5), 1368–1377.

Liu, Y., Whitman, W.B., 2008. Metabolic, phylogenetic, and ecological diversity of the methanogenic archaea. Annals of the new York Academy of Sciences 1125(1), 171–189.

Lovley, D.R., Klug, M.J., 1986. Model for the distribution of sulfate reduction and methanogenesis in freshwater sediments. Geochim. Cosmochim. Acta 50, 11–18.

Lovley, D.R., Phillips, E.J., 1987. Competitive mechanisms for inhibition of sulfate reduction and methane production in the zone of ferric iron reduction in sediments. Applied and Environmental Microbiology 53(11), 2636–2641.

López-Archilla, A.I., Moreira, D., Velasco, S., López-García, P., 2007. Archaeal and bacterial community composition of a pristine coastal aquifer in Donana National Park, Spain. Aquatic microbial ecology 47(2), 123–139.

Maltby, J., Sommer, S., Dale, A., Treude, T., 2016. Microbial methanogenesis in the sulfate-reducing zone of surface sediments traversing the Peruvian margin. Biogeosciences 13(1), 283–299. doi.org/10.5194/bg-13-283-2016.

Martin, M., 2011. Cutadapt removes adapter sequences from high-throughput sequencing reads. EMBnet. journal 17(1), 10–12.

Martinez-Cruz, K., Sepulveda-Jauregui, A., Casper, P., Anthony, K.W., Smemo, K.A., Thalasso, F., 2018. Ubiquitous and significant anaerobic oxidation of methane in freshwater lake sediments. Water research 144, 332–340.

Mausz, M.A., Chen, Y., 2019. Microbiology and ecology of methylated amine metabolism in marine ecosystems. Current Issues in Molecular Biology 33(1), 133–148.

McTaggart, T.L., Beck, D.A., Setboonsarng, U., Shapiro, N., Woyke, T., Lidstrom, M.E., Kalyuzhnaya, M.G., Chistoserdova, L., 2015. Genomics of methylotrophy in Gram-positive methylamine-utilizing bacteria. Microorganisms 3(1), 94–112.

Meyers, P.A., 1994. Preservation of elemental and isotopic source identification of sedimentary organic matter. Chemical geology 114(3-4), 289–302.

Michaelis, W., Seifert, R., Nauhaus, K., Treude, T., Thiel, V., Blumenberg, M., Knittel, K., Gieseke, A., Peterknecht, K., Pape, T., Boetius, A., Aman, A., Jørgensen, B.B., Widdel, F., Peckmann, J., Pimenov, N.V., Gulin, M., 2002. Microbial reefs in the Black Sea fueled by anaerobic oxidation of methane. Science 297, 1013–1015.

Moffett, K.B., Robinson, D.A., Gorelick, S.M., 2010. Relationship of salt marsh vegetation zonation to spatial patterns in soil moisture, salinity, and topography. Ecosystems 13, 1287–1302.

Montalto, F.A., Steenhuis, T.S., Parlange, J.-Y., 2006. The hydrology of Piermont Marsh, a reference for tidal marsh restoration in the Hudson river estuary, New York. Journal of Hydrology 316(1-4), 108–128.

Motelica-Heino, M., Naylor, C., Zhang, H., Davison, W., 2003. Simultaneous release of metals and sulfide in lacustrine sediment. Environmental Science & Technology 37(19), 4374–4381.

Murali, R., Yu, H., Speth, D.R., Wu, F., Metcalfe, K.S., Crémière, A., Laso-Pèrez, R., Malmstrom, R.R., Goudeau, D., Woyke, T., 2023. Physiological potential and evolutionary trajectories of syntrophic sulfate-reducing bacterial partners of anaerobic methanotrophic archaea. PLoS Biology 21(9), e3002292.

Newton, A., Icely, J., Cristina, S., Perillo, G.M., Turner, R.E., Ashan, D., Cragg, S., Luo, Y., Tu, C., Li, Y., 2020. Anthropogenic, direct pressures on coastal wetlands. Frontiers in Ecology and Evolution 8, 144.

Oksanen, J., Simpson, G., Blanchet, F., Kindt, R., Legendre, P., Minchin, P., O’Hara, R., Solymos, P., Stevens, M., Szoecs, E., 2022. vegan: Community Ecology Package. R package version 2.6-2. 2022. Google Scholar There is no corresponding record for this reference.

Oremland, R.S., Marsh, L.M., Polcin, S., 1982. Methane production and simultaneous sulphate reduction in anoxic, salt marsh sediments. Nature 296, 143–145.

Oremland, R.S., Polcin, S., 1982. Methanogenesis and sulfate reduction: competitive and noncompetitive substrates in estuarine sediments. Appl. Environ. Microbiol. 44(6), 1270–1276.

Oremland, R.S., Taylor, B.F., 1978. Sulfate reduction and methanogenesis in marine sediments. Geochimica et Cosmochimica Acta 42(2), 209–214.

Oren, A., 1990. Formation and breakdown of glycine betaine and trimethylamine in hypersaline environments. Antonie van Leeuwenhoek 58, 291–298.

Oren, A., 2011. Thermodynamic limits to microbial life at high salt concentrations. Environmental microbiology 13(8), 1908–1923.

Oren, A., 2014. The family methanoregulaceae. The prokaryotes: other major lineages of bacteria and the archaea, 253-258.

Orphan, V.J., House, C.H., Hinrichs, K.-U., McKeegan, K.D., De Long, E.F., 2001. Methane-consuming Archaea revealed by directly coupled isotopic and phylogenetic analysis. Science 293, 484–487.

Park, H.S., Lin, S., Voordouw, G., 2008. Ferric iron reduction by Desulfovibrio vulgaris Hildenborough wild type and energy metabolism mutants. Antonie Van Leeuwenhoek 93, 79–85.

Parkes, R.J., Brock, F., Banning, N., Hornibrook, E.R., Roussel, E.G., Weightman, A.J., Fry, J.C., 2012. Changes in methanogenic substrate utilization and communities with depth in a salt-marsh, creek sediment in southern England. Estuarine, Coastal and Shelf Science 96, 170–178.

Postma, D., Jakobsen, R., 1996. Redox zonation: equilibrium constraints on the Fe (III)/SO4-reduction interface. Geochimica et Cosmochimica Acta 60(17), 3169–3175.

Precht, E., Franke, U., Polerecky, L., Huettel, M., 2004. Oxygen dynamics in permeable sediments with wave-driven pore water exchange. Limnol. Oceanogr. 49(3), 693–705.

Quast, C., Pruesse, E., Yilmaz, P., Gerken, J., Schweer, T., Yarza, P., Peplies, J., Glöckner, F.O., 2012. The SILVA ribosomal RNA gene database project: improved data processing and web-based tools. Nucleic acids research 41(D1), D590–D596.

Ragsdale, S.W., Pierce, E., 2008. Acetogenesis and the Wood–Ljungdahl pathway of CO2 fixation. Biochimica et Biophysica Acta (BBA)-Proteins and Proteomics 1784(12), 1873–1898.

Reddy, K.R., DeLaune, R.D., 2008. Biogeochemistry of Wetlands. CRC Press, Taylor & Francis Group, Boca Raton.

Reddy, K.R., DeLaune, R.D., Inglett, P.W., 2022. Biogeochemistry of wetlands: science and applications. CRC press.

Reeburgh, W.S., 2007. Oceanic methane biogeochemistry. Chem. Rev. 107(2), 486–513.

Reyes, C., Schneider, D., Thürmer, A., Kulkarni, A., Lipka, M., Sztejrenszus, S.Y., Böttcher, M.E., Daniel, R., Friedrich, M.W., 2017. Potentially active iron, sulfur, and sulfate reducing bacteria in Skagerrak and Bothnian Bay sediments. Geomicrobiology Journal 34(10), 840–850.

Rocha, C., 1998. Rhythmic ammonium regeneration and flushing in intertidal sediments of the Sado estuary. Limnology and Oceanography 43(5), 823–831.

Rooze, J., Egger, M., Tsandev, I., Slomp, C.P., 2016. Iron-dependent anaerobic oxidation of methane in coastal surface sediments: Potential controls and impact. Limnology and Oceanography 61(S1), S267–S282.

Ruff, S.E., Kuhfuss, H., Wegener, G., Lott, C., Ramette, A., Wiedling, J., Knittel, K., Weber, M., 2016. Methane seep in shallow-water permeable sediment harbors high diversity of anaerobic methanotrophic communities, Elba, Italy. Frontiers in Microbiology 7, 374.

Rusch, A., Forster, S., Huettel, M., 2001. Bacteria, diatoms and detritus in an intertidal sandflat subject to advective transort across the water-sediment interface. Biogeochemistry 55, 1–27.

Saunois, M., Stavert, A.R., Poulter, B., Bousquet, P., Canadell, J.G., Jackson, R.B., Raymond, P.A., Dlugokencky, E.J., Houweling, S., Patra, P.K., 2020. The global methane budget 2000–2017. Earth system science data 12(3), 1561–1623.

Scheller, S., Yu, H., Ghadwick, G.L., McGlynn, S.E., Orphan, V.J., 2016. Artificial electron acceptors decouple archaeal methane oxidation from sulfate reduction. Science 351, 703–707.

Schnakenberg, A., Aromokeye, D.A., Kulkarni, A., Maier, L., Wunder, L.C., Richter-Heitmann, T., Pape, T., Ristova, P.P., Bühring, S.I., Dohrmann, I., 2021. Electron acceptor availability shapes anaerobically methane oxidizing archaea (ANME) communities in South Georgia sediments. Frontiers in Microbiology 12, 617280.

Segarra, K.E., Comerford, C., Slaughter, J., Joye, S.B., 2013. Impact of electron acceptor availability on the anaerobic oxidation of methane in coastal freshwater and brackish wetland sediments. Geochimica et Cosmochimica Acta 115, 15–30.

Smith, M.W., Davis, R.E., Youngblut, N.D., Kärnä, T., Herfort, L., Whitaker, R.J., Metcalf, W.W., Tebo, B.M., Baptista, A.M., Simon, H.M., 2015. Metagenomic evidence for reciprocal particle exchange between the mainstem estuary and lateral bay sediments of the lower Columbia River. Frontiers in Microbiology 6, 1074.

Smith, M.W., Herfort, L., Rivers, A.R., Simon, H.M., 2019. Genomic signatures for sedimentary microbial utilization of phytoplankton detritus in a fast-flowing estuary. Frontiers in Microbiology 10, 2475.

Sorokin, D.Y., Merkel, A.Y., Abbas, B., Makarova, K.S., Rijpstra, W.I.C., Koenen, M., Sinninghe Damsté, J.S., Galinski, E.A., Koonin, E.V., Van Loosdrecht, M.C., 2018. Methanonatronarchaeum thermophilum gen. nov., sp. nov. and’Candidatus Methanohalarchaeum thermophilum’, extremely halo (natrono) philic methyl-reducing methanogens from hypersaline lakes comprising a new euryarchaeal class Methanonatronarchaeia classis nov. International journal of systematic and evolutionary microbiology 68(7), 2199–2208.

Sun, J., Evans, P.N., Gagen, E.J., Woodcroft, B.J., Hedlund, B.P., Woyke, T., Hugenholtz, P., Rinke, C., 2021. Recoding of stop codons expands the metabolic potential of two novel Asgardarchaeota lineages. ISME communications 1(1), 30.

Taubert, M., Grob, C., Howat, A.M., Burns, O.J., Pratscher, J., Jehmlich, N., von Bergen, M., Richnow, H.H., Chen, Y., Murrell, J.C., 2017. Methylamine as a nitrogen source for microorganisms from a coastal marine environment. Environmental microbiology 19(6), 2246–2257.

Thauer, R.K., 1998. Biochemistry of methanogenesis: a tribute to Marjory Stephenson. Microbiology 144, 2377–2406.

Thureborn, P., Franzetti, A., Lundin, D., Sjöling, S., 2016. Reconstructing ecosystem functions of the active microbial community of the Baltic Sea oxygen depleted sediments. PeerJ 4, e1593.

Tilbrook, B.D., Karl, D.M., 1995. Methane sources, distributions and sinks from California coastal waters to the oligotrophic North Pacific gyre. Marine Chemistry 49(1), 51–64.

Timmers, P.H.A., Welte, C.U., Koehorst, J.J., Plugge, C.M., Jetten, M.S.M., Stams, A.J.M., 2017. Reverse methanogenesis and respiration in methanotrophic archaea. Archaea Article ID 1654237.

Treude, T., Krüger, M., Boetius, A., Jørgensen, B.B., 2005. Environmental control on anaerobic oxidation of methane in the gassy sediments of Eckernförde Bay (German Baltic). Limnol. Oceanogr. 50, 1771–1786.

Valenzuela, E.I., Avendaño, K.A., Balagurusamy, N., Arriaga, S., Nieto-Delgado, C., Thalasso, F., Cervantes, F.J., 2019. Electron shuttling mediated by humic substances fuels anaerobic methane oxidation and carbon burial in wetland sediments. Science of The Total Environment 650, 2674–2684.

Vizza, C., West, W.E., Jones, S.E., Hart, J.A., Lamberti, G.A., 2017. Regulators of coastal wetland methane production and responses to simulated global change. Biogeosciences. 14 (2): 431–446. 14(2), 431–446.

Waite, D.W., Chuvochina, M., Pelikan, C., Parks, D.H., Yilmaz, P., Wagner, M., Loy, A., Naganuma, T., Nakai, R., Whitman, W.B., 2020. Proposal to reclassify the proteobacterial classes Deltaproteobacteria and Oligoflexia, and the phylum Thermodesulfobacteria into four phyla reflecting major functional capabilities. International Journal of Systematic and Evolutionary Microbiology 70(11), 5972–6016.

Wallenius, A.J., Dalcin Martins, P., Slomp, C.P., Jetten, M.S., 2021. Anthropogenic and environmental constraints on the microbial methane cycle in coastal sediments. Frontiers in Microbiology 12, 631621.

Wang, X.-c., Lee, C., 1990. The distribution and adsorption behavior of aliphatic amines in marine and lacustrine sediments. Geochimica et Cosmochimica Acta 54(10), 2759–2774.

Wang, X.-C., Lee, C., 1993. Adsorption and desorption of aliphatic amines, amino acids and acetate by clay minerals and marine sediments. Marine Chemistry 44(1), 1–23.

Wang, X.-C., Lee, C., 1994. Sources and distribution of aliphatic amines in salt marsh sediment. Organic Geochemistry 22(6), 1005–1021.

Wang, Y., He, L., Liu, J., Arndt, K.A., Mazza Rodrigues, J.L., Zona, D., Lipson, D.A., Oechel, W.C., Ricciuto, D.M., Wullschleger, S.D., 2024. Intensified Positive Arctic–Methane Feedback under IPCC Climate Scenarios in the 21st Century. Ecosystem Health and Sustainability 10, 0185.

Weber, K.A., Achenbach, L.A., Coates, J.D., 2006. Microorganisms pumping iron: anaerobic microbial iron oxidation and reduction. Nature Reviews Microbiology 4(10), 752–764.

Wehrmann, L.M., Risgaard-Petersen, N., Schrum, H.N., Walsh, E.A., Huh, Y., Ikehara, M., Pierre, C., D’Hondt, S., Ferdelman, T.G., Ravelo, A.C., 2011. Coupled organic and inorganic carbon cycling in the deep subseafloor sediment of the northeastern Bering Sea Slope (IODP Exp. 323). Chemical Geology 284(3-4), 251–261.

Winfrey, M.R., Ward, D.M., 1983. Substrates for sulfate reduction and methane p roduction in i ntertidal sediments. Appl. Environm. Microbiol 45(1), 193–199.

Xia, S., Song, Z., Li, Q., Guo, L., Yu, C., Singh, B.P., Fu, X., Chen, C., Wang, Y., Wang, H., 2021. Distribution, sources, and decomposition of soil organic matter along a salinity gradient in estuarine wetlands characterized by C: N ratio, δ13C-δ15N, and lignin biomarker. Global Change Biology 27(2), 417–434.

Xiao, K., Beulig, F., Roy, H., Jorgensen, B., Risgaard-Petersen, N., 2018. Methylotrophic methanogenesis fuels cryptic methane cycling in marine surface sediment. Limnology and Oceanography 63(4), 1519–1527. doi.org/10.1002/lno.10788.

Xiao, K.-Q., Beulig, F., Kjeldsen, K.U., Jørgensen, B.B., Risgaard-Petersen, N., 2017. Concurrent methane production and oxidation in surface sediment from Aarhus Bay, Denmark. Frontiers Microbiol. 8, doi: 10.3389/fmicb.2017.01198.

Xiao, K.-Q., Moore, O.W., Babakhani, P., Curti, L., Peacock, C.L., 2022. Mineralogical control on methylotrophic methanogenesis and implications for cryptic methane cycling in marine surface sediment. Nature Communications 13(1), 1–9.

Yan, Z., Du, K., Yan, Y., Huang, R., Zhu, F., Yuan, X., Wang, S., Ferry, J.G., 2023. Respiration-driven methanotrophic growth of diverse marine methanogens. Proceedings of the National Academy of Sciences 120(39), e2303179120.

Yan, Z., Joshi, P., Gorski, C.A., Ferry, J.G., 2018. A biochemical framework for anaerobic oxidation of methane driven by Fe (III)-dependent respiration. Nature communications 9(1), 1642.

Young, C.M., 2003. Reproduction, development and life-history traits, in: Tyler, P.A. (Ed.) Deep-sea ecosystems. Elsevier, Amsterdam.

Yuan, J., Ding, W., Liu, D., Xiang, J., Lin, Y., 2014. Methane production potential and methanogenic archaea community dynamics along the Spartina alterniflora invasion chronosequence in a coastal salt marsh. Applied microbiology and biotechnology 98, 1817–1829.

Yuan, J., Liu, D., Ji, Y., Xiang, J., Lin, Y., Wu, M., Ding, W., 2019. Spartina alterniflora invasion drastically increases methane production potential by shifting methanogenesis from hydrogenotrophic to methylotrophic pathway in a coastal marsh. Journal of Ecology 107(5), 2436–2450.

Zemskaya, T., Bukin, S., Lomakina, A., Pavlova, O., 2021. Microorganisms in the sediments of Lake Baikal, the deepest and oldest lake in the world. Microbiology 90(3), 298–313.

Zhao, M., Han, G., Li, J., Song, W., Qu, W., Eller, F., Wang, J., Jiang, C., 2020. Responses of soil CO2 and CH4 emissions to changing water table level in a coastal wetland. Journal of Cleaner Production 269, 122316.

Zhuang, G.-C., Elling, F.J., Nigro, L.M., Samarkin, V., Joye, S.B., Teske, A., Hinrichs, K.-U., 2016. Multiple evidence for methylotrophic methanogenesis as the dominant methanogenic pathway in hypersaline sediments from the Orca Basin, Gulf of Mexico. Geochim. Cosmochim. Acta 187, 1–20.

Zhuang, G.-C., Heuer, V.B., Lazar, C.S., Goldhammer, T., Wendt, J., Samarkin, V.A., Elvert, M., Teske, A.P., Joye, S.B., Hinrichs, K.-U., 2018. Relative importance of methylotrophic methanogenesis in sediments of the Western Mediterranean Sea. Geochim. Cosmochim. Acta 224.

Zhuang, G.-C., Lin, Y.-S., Bowles, M.W., Heuer, V.B., Lever, M.A., Elvert, M., Hinrichs, K.- U., 2017. Distribution and isotopic composition of trimethylamine, dimethylsulfide and dimethylsulfoniopropionate in marine sediments. Mar. Chem. 196, 35–46.

Zhuang, G.-C., Montgomery, A., Joye, S.B., 2019. Heterotrophic metabolism of C1 and C2 low molecular weight compounds in northern Gulf of Mexico sediments: Controlling factors and implications for organic carbon degradation. Geochimica et Cosmochimica Acta 247, 243–260.

Ziebis, W., Huettel, M., Forster, S., 1996. The impact of biogenic sediment topography on oxygen fluxes in permeable sediments. Mar. Ecol. Prog. Ser. 140, 227–237.

